# Analyzing Presymptomatic Tissue to Gain Insights into the Molecular and Mechanistic Origins of Late-Onset Degenerative Trinucleotide Repeat Disease

**DOI:** 10.1101/2020.02.25.964403

**Authors:** Yongjun Chu, Jiaxin Hu, Hanquan Liang, Mohammed Kanchwala, Chao Xing, Walter Beebe, Charles B. Bowman, Xin Gong, David R. Corey, V. Vinod Mootha

**Author notes:** Corresponding authors: V. Vinod Mootha M.D., Department of Ophthalmology, Eugene McDermott Center for Human Growth and Development, 5323 Harry Hines Blvd., Dallas, TX 75390-9057, David R. Corey Ph.D., Department of Pharmacology, Department of Biochemistry, 5323 Harry Hines Blvd., Dallas, TX 75390.

## Abstract

How genetic defects trigger the molecular changes that cause late-onset disease is important for understanding disease progression and therapeutic development. Fuchs’ endothelial corneal dystrophy (FECD) is an RNA-mediated disease caused by a trinucleotide CUG expansion in an intron within the *TCF4* gene. The mutant intronic CUG RNA is present at 1-2 copies per cell, posing a challenge to understand how a rare RNA can cause disease. Late-onset FECD is a uniquely advantageous model for studying how RNA triggers disease because; 1) Affected tissue is routinely removed during surgery; 2) The expanded CUG mutation is one of the most prevalent disease-causing mutations, making it possible to obtain pre-symptomatic tissue from eye bank donors to probe how gene expression changes precede disease; and 3) The affected tissue is a homogeneous single cell monolayer, facilitating accurate transcriptome analysis. Here we use RNA sequencing (RNAseq) to compare tissue from individuals who are pre-symptomatic (Pre_S) to tissue from patients with late stage FECD (FECD_REP). The abundance of mutant repeat intronic RNA in Pre_S and FECD_REP tissue is elevated due to increased half-life in a corneal cell-specific manner. In Pre_S tissue, changes in splicing and extracellular matrix gene expression foreshadow the changes observed in advanced disease and predict the activation of the fibrosis pathway and immune system seen in late-stage patients. The absolute magnitude of splicing changes is similar in presymptomatic and late stage tissue. Our data identify gene candidates for early drivers of disease and biomarkers that may represent diagnostic and therapeutic targets for FECD. We conclude that changes in alternative splicing and gene expression are observable decades prior to the diagnosis of late-onset trinucleotide repeat disease.

## Introduction

An inherent paradox of inherited late-onset degenerative disease is the fact that the genetic basis for disease exists years before symptoms arise. At some point the genetic defect begins to trigger a cascade of cellular changes that result in disease, but how is the cascade initiated? Unfortunately, the search for early drivers of many genetic diseases that involve mutant RNA is hindered by lack of accessible pre-symptomatic tissue to identify early changes in gene expression that occur years prior to diagnosis. Corneal tissue provides an accessible source of both pre- and post-symptomatic tissue to explore the molecular origins of inherited late-onset disease.

Corneal diseases represent one of the leading causes of vision loss and blindness globally (1). Inherited corneal dystrophies can compromise the structure and transparency of the cornea. Late-onset Fuchs’ endothelial corneal dystrophy (FECD) is one of the most common genetic disorders, affecting four percent of the population in the United States over the age of forty (2–4). The corneal endothelium is the inner hexagonal monolayer responsible for maintenance of stromal dehydration and corneal clarity.

In FECD, the post-mitotic endothelium undergoes premature senescence and apoptosis (5–11). Descemet’s membrane, the basement membrane of the endothelium, becomes diffusely thickened and develops focal excrescences called guttae. Guttae are clinically diagnostic of FECD by slit-lamp biomicroscopy (12). Confluence of central guttae and concomitant loss of endothelial cell density results in corneal edema, scarring, and loss of vision, making FECD the leading indication for corneal transplantation in the United States and developed world (13, 14).

Two thirds of FECD cases are caused by an expansion of the trinucleotide CUG within the *TCF4* gene (15–18) making the corneal dystrophy the most common human disorder mediated by simple DNA repeats. FECD can also be caused by a CUG expansion within the 3’-untranslated region (3’-UTR) of the *DMPK* gene (19–21), implicating mutant expanded CUG RNA as the root cause for repeat-associated FECD (FECD_REP). The remaining one third of patients lack the expanded repeat (FECD_NR) but the two types of FECD are indistinguishable during normal clinical observation.

FECD can be treated with corneal transplantation and surgical outcomes have improved following the development of endothelial keratoplasty in which a single-cell endothelium monolayer is transplanted using donor corneal tissue (22). While effective, this approach is limited by the availability of donor corneas and access to suitable treatment centers. A deeper understanding of FECD might permit the development of pharmacological approaches that might reach a larger patient population and reduce or delay the need for surgical intervention. Successful drug development would benefit from a more detailed understanding of disease progression and the fundamental drivers of early disease.

Trinucleotide or hexanucleotide repeats are the cause for many delayed-onset neurodegenerative disorders such as myotonic dystrophy type 1 and *C9orf72* amyotrophic lateral sclerosis/frontal temporal dementia (ALS/FTD) (23). FECD has unique advantages as a model for gaining insight into the molecular mechanisms of delayed-onset degenerative disease; 1) The tissue is more readily available for analysis because it is routinely removed during surgery (**Figure 1A-C**); 2) Samples are a single near homogeneous layer of cells, facilitating RNAseq and other methods for analyzing gene expression (**Figure 1D**); 3) The corneal endothelium and disease progression can be evaluated visually during a standard clinical examination; 4) Like peripheral neurons, the corneal endothelium originates from neural crest and may serve as a model post-mitotic tissue to examine the effects of toxic repeat RNA over time; 5) Donor corneal tissues that are not used for surgery are available from eye bank samples for comparison; 6) Because the prevalence of the triplet repeat mutation within the *TCF4* gene is relatively high, donor eyes provide a significant number of pre-symptomatic samples (Pre_S) that possess the CUG expanded repeat; 7) The availability of tissue from four different cohorts (**Figure 2**), control, Pre_S, FECD_NR, FECD_REP allows multiple cross-comparisons into the different stages and types of FECD. Analysis of the Pre_S cohort has the potential to gain insights into early drivers of disease.

**Figure 1.**
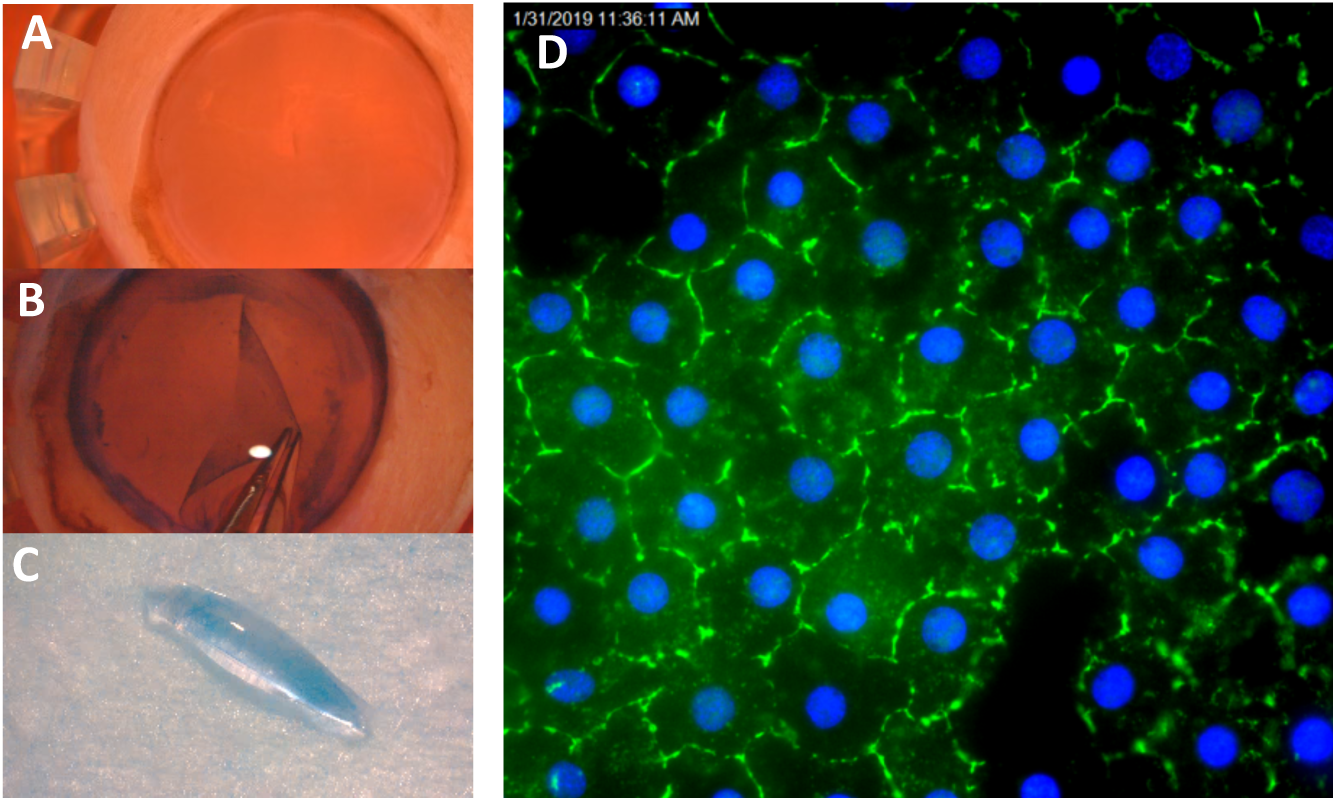
Preparation of homogeneous tissue monolayers for analysis. **(A)** Human donor cornea in corneal viewing chamber with Optisol corneal storage media. **(B)** Corneal endothelium / Descemet’s membrane monolayer being dissected from underlying stromal tissue. **(C)** Monolayer of corneal endothelial cells assumes a “scroll” shape. This single cell monolayer will be used for sequencing and other analyses. **(D)** Immunostaining of monolayer of cells with corneal endothelial specific marker, zonula occudens-1 (ZO-1) (Blue-DAPI).

**Figure 2.**
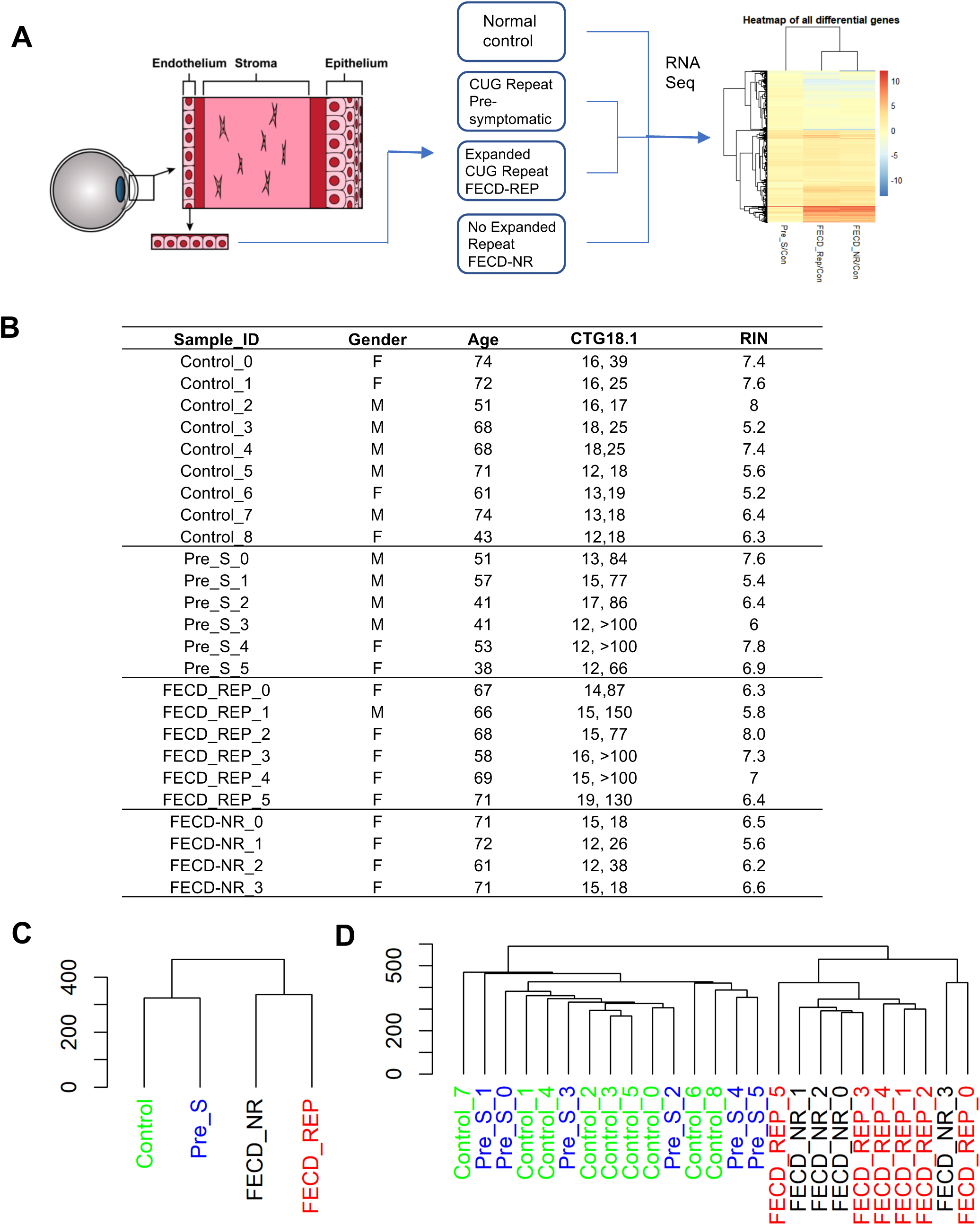
Experimental scheme and sample cohorts. (**A**) Homogenous samples of corneal endothelium are surgically removed from normal, presymptomatic and, affected (FECD or FECD_NR) tissue bank donors or patients and used for RNAseq followed by data analysis. (**B**) Detailed description of tissue samples. (**C**) Dendrogram for four groups of samples and (**D**) all replicates. RIN:RNA integrity number. CTG18.1: the number of repeats on each chromosome.

Here, we use the advantages gained from analyzing the corneal endothelial monolayer to evaluate transcriptomic data from four cohorts of corneal endothelial tissue: control, Pre_S, FECD_REP, and FECD_NR. We identify extensive and large magnitude changes in RNA splicing that are shared between Pre_S and FECD-REP cohorts but are not observed in FECD_NR samples. Levels of gene expression are changed in Pre_S samples relative to controls and suggest early triggering of the fibrosis pathway prior to clinical observation of disease symptoms. In late stage disease, pathways related to fibrosis and activation of immune system are shared by FECD_REP and FECD_NR but mitochondrial dysfunction is more pronounced in the FECD_REP cohort. These results lay a basis for understanding the onset of FECD and other trinucleotide repeat diseases and provide potential targets for therapeutic design.

## Materials and Methods

### Isolation of Corneal Tissue

The study was conducted in compliance with the tenets of the Declaration of Helsinki and with the approval of the institutional review board of the University of Texas Southwestern Medical Center (UTSW). Subjects underwent a complete eye examination including slit lamp biomicroscopy by a cornea fellowship-trained ophthalmologist. Patients underwent endothelial keratoplasty for FECD severity Krachmer grade 5 (≧5 mm central confluent guttae without stromal edema) or 6 (≧5 mm central confluent guttae with stromal edema) assessed by slit lamp microscopy (56). After surgery, surgically explanted endothelium-Descemet’s membrane monolayers were preserved in Optisol GS corneal storage media (Bausch & Lomb, Rochester, NY) prior to storage at −80 Celsius. Genomic DNA was extracted from peripheral blood leukocytes of each study subject using Autogen Flexigene (Qiagen, Valencia, CA).

Corneal endothelial samples from post-mortem donor corneas preserved in Optisol GS corneal storage media (Bausch & Lomb, Rochester, NY) were obtained from the eye bank of Transplant Services at UT Southwestern. Certified eye bank technicians screened the donor corneal endothelium with slit lamp biomicroscopy and Cellchek EB-10 specular microscopy (Konan Medical). Endothelium-Descemet’s membrane monolayers from donor corneas were micro-dissected and stored as previously described (43). DNA from the remaining corneal tissue of each sample was extracted with TRIzol reagent (ThermoScientific).

### TCF4 CTG18.1 polymorphism genotyping

Genomic DNA from subjects’ peripheral leukocytes or corneal tissue was used for genotyping. The CTG18.1 trinucleotide repeat polymorphism in the *TCF4* gene was genotyped using a combination of short tandem repeat (STR) and triplet repeat primed polymerase chain reaction (TP-PCR) assays as we have previously described (20). For the STR assay, a pair of primers flanking the CTG18.1 locus was utilized for PCR amplification with one primer labeled with FAM on 5′ end. The TP-PCR assay was performed using the 5’ FAM labeled primer specific for the repeat locus paired with repeat sequence targeted primers for PCR amplification. PCR amplicons were loaded on an ABI 3730XL DNA analyzer (Applied Biosystems, Foster City, CA, USA) and the results analyzed with ABI GeneMapper 4.0 (Applied Biosystems). Large triplet repeat expansions were sized by Southern blot analysis using digoxigenin labeled probes.

### RNA Isolation and Sequencing

Total RNA was isolated from each of 25 tissue samples (6 FECD_REP, 4 FECD_NR, 6 Pre_S and 9 Controls) by homogenization in QIAzol lysis reagent, chloroform extraction and isolation with NucleoSpin RNA XS (Macherey-Nagel GmbH & Co., Germany). RNA quantity and quality were determined by Bioanalyzer 2100. RNA libraries were prepared for each tissue sample with high RIN (> 5.0), using the TruSeq RNA sample Prep kit version 2 (Illumina, San Diego, CA, USA). For TruSeq stranded total RNAseq, ribosomal transcripts were depleted from total RNA, using Ribo-Zero Gold RNA removal kit followed by replacement of deoxythymidine triphosphate (dTTP) with deoxyuridine triphosphate (dUTP) during reverse transcription in the second strand synthesis, using TruSeq stranded total library preparation kit. The resulting libraries were minimally amplified to enrich for fragments using adapters on both ends and then quantified for sequencing at eight samples/flow cell by using a NextSeq 500/550 (Illumina) sequencer (PE 150).

### Analysis of Differentially Expressed or Spliced Genes

Whole transcriptomic sequencing data from each tissue sample was analyzed using an analysis pipeline which includes STAR for initial mapping and Cufflinks (v2.21) for gene and isoform differential analysis, among other publicly available programs. For gene/isoform differential analysis, the minimum expression level of 1.5 FPKM and an FDR < 0.05 were chosen as the threshold. The meta gene pathway analysis was carried out with IPA (Qiagene). The binary alignment map files from STAR were analyzed using rMATS (v.4.0) that quantitates the expression level of alternatively spliced genes between groups. To find the most significant events, we used stringent filtering criteria within rMATS to perform pairwise comparisons among 4 groups: percentage of spliced in (PSI) changes > 0.15; FDR < 0.001. For PSI, rMATS calculates a value for every differential splicing event, providing a range from 0 to 1, with 0 being completely excluded and 1 being uniformly included in the splicing products. Alternative splicing events were also compared to those obtained in tibialis anterior muscle of myotonic dystrophy type one patients (DM1). DM1 raw data was obtained from Gene Expression Omnibus (GSE86356, 6 Control and 6 DM1 tissue samples were used) and analyzed similarly as the FECD data (visit DMseq.org for more information).

### Measurement of TCF4 RNA half-life

F35T corneal endothelial cell line and a control corneal endothelial cells (W4056-17-001579) were cultured as described (44) The C9 and VVM84 skin fibroblasts were maintained at 37°C and 5% CO_2_ in Minimal Essential Media Eagle (MEM) (Sigma, M4655) supplemented with 15% heat inactivated fetal bovine serum (Sigma) and 0.5% MEM nonessential amino acids (Sigma).

Endothelial or fibroblast cells were seeded in 6-well plate at 90% confluence. At the next day, actinomycin D was added into the wells at 5 µg/mL final concentration. Cells were harvest using Trizol agent (Sigma) at different time point. The *TCF4* RNA levels were analyzed by qPCR.

### Validation of Differential Gene Expression and Alternative Splicing Patterns

Total RNA was extracted from control, Pre_S or FECD corneal endothelial tissues (**Supplemental Table 1**) by NucleoSpin RNA XS kit (Macherey-Nagel). cDNAs were prepared by reverse transcription. RT-PCR was performed using ChoiceTaq Blue Mastermix (Denville). PCR amplification was as follow: 94 °C for 3 minutes (1 cycle), 94 °C denaturation for 30 sec, 60 °C annealing for 30 sec, and 72 °C extension for 1 min (38 cycles), and a 7 minute 72 °C extension. The PCR primers were used as reported (28) (**Supplemental Table 2**). The amplification products were separated by 1.5% agarose gel electrophoresis. *q*PCR experiments were performed on a 7500 real-time PCR system (Applied Biosystems) using iTaq SYBR Green Supermix (Bio-Rad). Data was normalized relative to levels of *RPL19* mRNA.

## Results

### Experimental Design

Samples were homogeneous endothelial cell monolayers surgically removed from patients or dissected from donor tissue (**Figure 1, Figure 2A**). The homogenous nature of the tissue samples, in contrast to the more complex mixtures of cells often contained in tissues related to other disease, facilitates subsequent analysis and data interpretation.

We compared four groups of human tissue by RNA sequencing (RNAseq) with attention to changes in alternative splicing, differential gene expression, and pathway analysis (**Figure 2A**). Tissue that was mutant for the CUG expanded repeat within intron 2 of the *TCF4* gene was obtained from pre-symptomatic eye bank donors (Pre_S) and from FECD patients after transplant surgery (FECD_REP). Tissue that was negative for the expanded CUG mutation within *TCF4* was obtained from eye bank donors (Control) and from patients with non-CUG related FECD (FECD_NR).

Control tissues from the eye bank were chosen for analysis that possessed normal endothelial morphology by specular microscopy, were negative for the expanded CUG mutation, and were from donors with ages comparable to our FECD patient cohort (**Figure 2B**). Obtaining tissue from Pre_S individuals was possible because of the relatively high prevalence, 3%, of the expanded triplet repeat mutation in *TCF4* gene within the general Caucasian population (2, 3). Pre_S tissue was identified by the presence of the CUG repeat expansion by genotyping. Normal corneal endothelial morphology (absence of central corneal guttae) in Pre_S tissues was confirmed by specular microscopy.

RNA sequencing (RNAseq) was performed on tissue samples from nine non-FECD/non-expanded repeat donors (Control), six pre-symptomatic donors with the expanded repeat (Pre_S), six patients with the expanded repeat and FECD (FECD_REP), and four patients with FECD who did not have the expanded CUG repeat mutation (FECD_NR) (**Figure 2B**). Only samples with RNA integrity numbers greater than 5 were used. We carried out a paired-end 150 nt RNA-Seq on Illumina NextSeq sequencer. We regularly obtained an output of 50-60 million raw paired reads per sample.

Clustering analysis of overall gene expression patterns revealed that samples from control and Pre_S donors were closer to one another than to samples from patients with FECD_REP or FECD_NR late stage disease (**Figure 2CD, Supplemental Figure 1AB**). This result is consistent with the clinical observation of large visible differences between diseased and non-diseased tissue and that the clinical manifestations of FECD_REP and FECD_NR are almost identical.

### Stability of TCF4 intron 2 in FECD mutant and non-mutant tissue

Analysis of RNAseq data from intron 2 (the intron that contains the CUG repeat) within the *TCF4* gene showed a striking difference between representative samples from individuals with the expanded CUG repeat mutation (FECD_REP and Pre_S) and individuals who lack the mutation (control and FECD_NR) (**Figure 3A**). Samples from the cohort of control individuals showed similar low numbers of reads upstream or downstream relative to the trinucleotide repeat region. RNA obtained from FECD_NR patients’ tissue also showed the same similarity between upstream and downstream reads.

**Figure 3.**
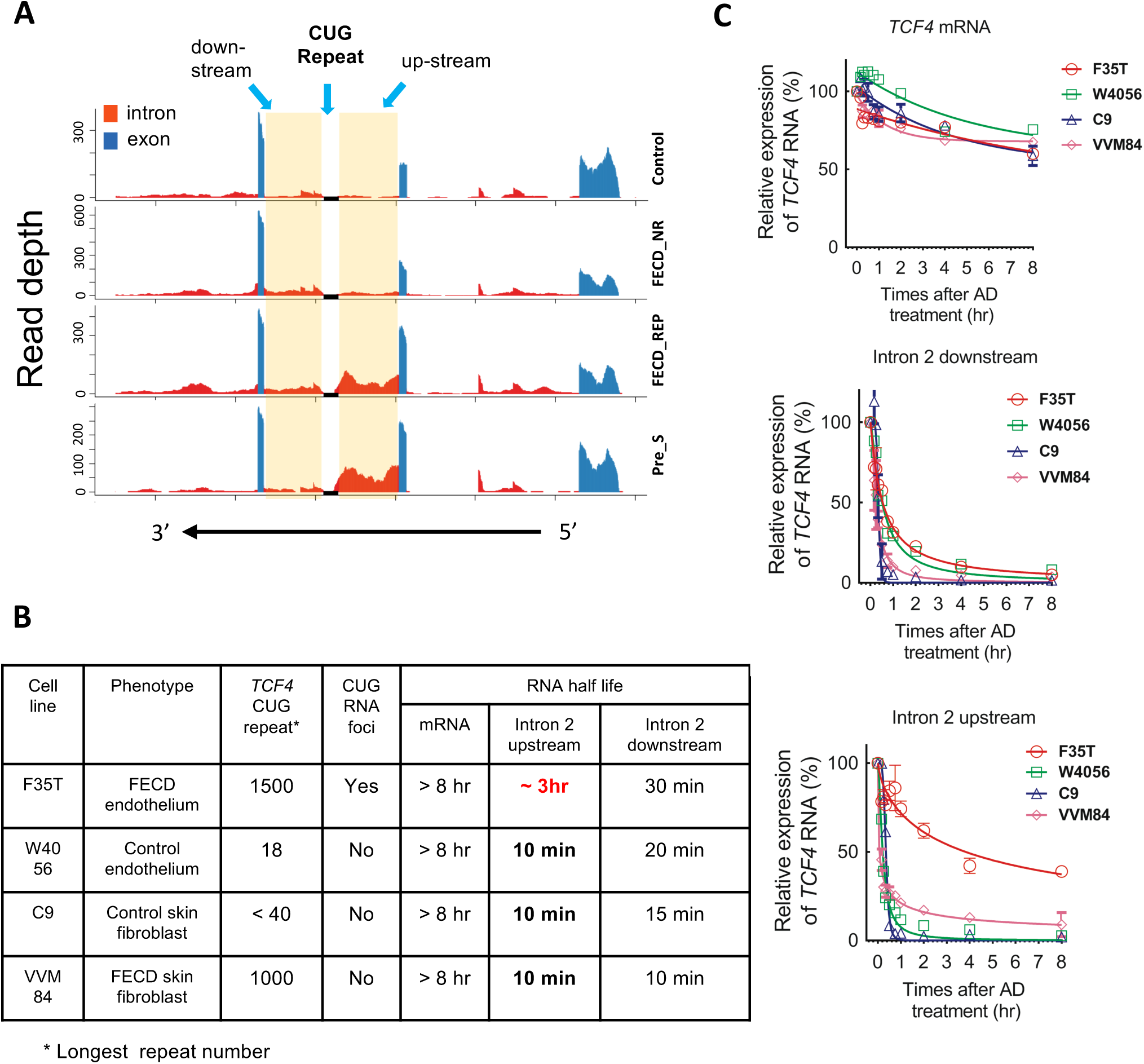
Intronic RNA is stabilized in Pre_S and FECD_REP tissue and FECD_REP patient-derived F35T corneal endothelial cells. (**A**) Representative RNAseq data showing relative read number for *TCF4* intron 2 RNA from each sample cohort. (**B**) Summary of RNA half-life of different cell lines. The much longer half life of intron 2 up-stream RNA in F35T cells is highlighted in red. (**C**) Graphs of time-dependent RNA decay following the actinomycin D treatment.

By contrast, both Pre_S and FECD_REP tissue showed more reads for intronic RNA upstream of the trinucleotide repeat relative to downstream (**Figure 3A**). The overall expression of mature *TCF4* mRNA from the four cohorts was not significantly different making haploinsufficiency of gene product less likely as the mechanism of disease (**Supplemental Figure 2**). These results suggest that an early molecular disease trigger – increased stability the mutant *TCF4* intron 2 upstream of the expanded repeat – occurs at the presymptomatic stage and distinguishes FECD_REP from FECD_NR in late stage disease.

To investigate factors that might contribute to the prevalence of upstream intronic reads, we examined cultured cells derived from FECD_REP patient corneal endothelium (F35T), FECD_REP patient skin fibroblasts (VVM84), control (without CUG expanded repeat) corneal endothelium (W4056), or control skin fibroblasts (C9) (**Figure 3BC**). VVM84 FECD skin fibroblasts and F35T FECD corneal endothelial cells both have the expanded CUG repeat, but the F35T corneal cells have nuclear CUG foci that can be detected by RNA-FISH while VVM84 skin cells lack detectable foci indicating cell-specificity for expanded CUG repeat RNA accumulation.

We treated cells with actinomycin D to arrest transcription and examine the half-life of mature *TCF4* mRNA and sequences either upstream our downstream relative to the intronic *TCF4* CUG repeat using quantitative PCR (qPCR) (**Figure 3BC**). Regardless of whether the expanded repeat mutation was present, the half-life of the mature *TCF4* mRNA was similar, >8 hours (**Figure 3C top**). Likewise, the half-life of the intron 2 downstream region was also similar in each cell line, varying from 10 to 30 minutes (**Figure 3C middle**). These data suggest that the mutation does not affect stability of the parent mRNA and has only a modest effect on the downstream region of intron 2.

By contrast, we observe a striking ∼20-fold increase in the half-life of the upstream region of intron 2 in F35T corneal cells that possess the expanded CUG mutation and have detectable foci (**Figure 3C bottom**). For cell lines that lacked expanded CUG nuclear foci, this region had a half-life of only 10 minutes. For F35T corneal cells, the half-life was three hours.

This increased half-life of upstream intronic RNA is consistent with the observation from RNAseq of many more reads covering the upstream region of intron 2 *TCF4* from Pre_S or FECD_REP tissue relative to samples from individuals in the control or FECD_NR cohorts who lack the expanded CUG repeat (**Figure 3A**). Stabilization of the upstream portion of *TCF4* intron 2 in corneal endothelial tissue and cultured corneal endothelial cells makes more mutant repeat RNA available to perturb gene expression.

### Changes in Alternative splicing triggered by the expanded repeat within TCF4

CUG repeat RNA is known to bind the splicing factors muscleblind-like 1 and 2 (MBNL1 and MNBL2) (24–27). Previous studies have proposed that sequestration of MBNL1 and MBNL2 reduces levels of available MBNL protein, causing the changes of splicing observed in tissue from FECD_REP patients (28–30) and in patients with myotonic dystrophy who possess expanded CUG repeats within 3’-untranslated region of the *DMPK* gene (24,25,31).

We used RNAseq to evaluate splicing changes between control tissue and the Pre_S, FECD_REP, and FECD_NR tissue cohorts (**Figure 4**). To classify changes, we used the FDR (false discovery rate) and delta PSI (the net change of inclusion percentage) as the determinant metrics. Any changes that are less than 0.001 on FDR and more than or equal to 0.15 on PSI were determined to be significant.

**Figure 4.**
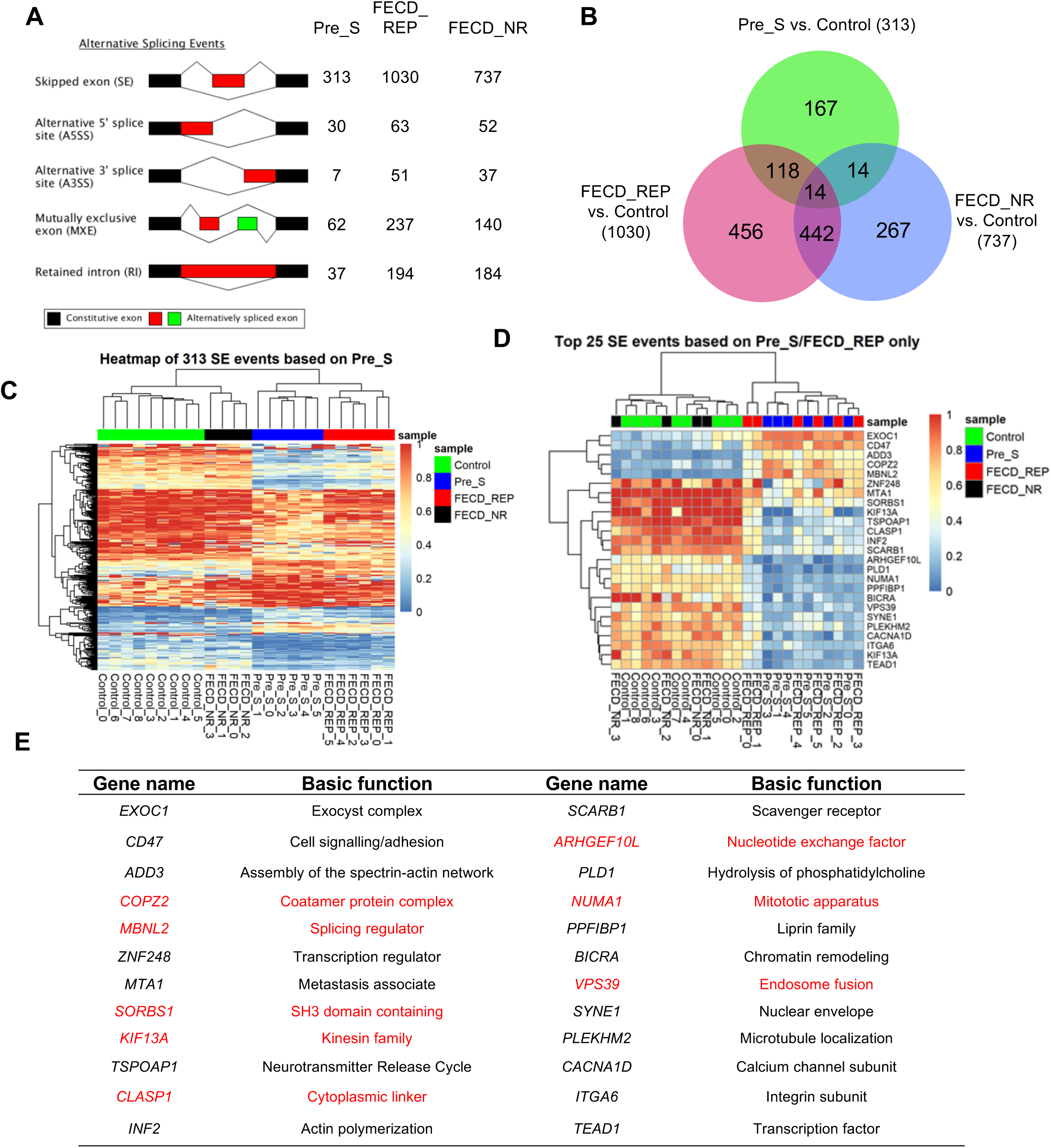
RNAseq data demonstrates changes in splicing in pre-symptomatic, FECD_REP, and FECD_NR tissue. **A**) Alternative splicing events in Pre_S, FECD_REP, and FECD_NR cohorts. (**B**) Overlap between sample cohorts for skipped exon (SE) events showing that late stage disease FECD_REP and Pre_S cohorts cluster differently from control or FECD_NR cohorts. (**C**) Heat map comparing inclusion levels of exons among 4 cohorts. The chosen SE events were based on the 313 significant skipped exon events identified in Pre_S tissues. The Pre_S cohort is more similar to FECD_REP than it is to the other two cohorts. (**D**) The similarity of Pre_S and FECD_REP is emphasized by a heat map comparing top 25 SE events in common between FECD_REP and Pre_S tissues and corresponding changes in FECD_NR and control tissues. (**E**) Identity of top 25 SE events in common between FECD_REP and Pre_S. Genes in red are also observed in RNAseq data from DM1 tissue, demonstrating substantial overlap despite different tissue origins. The threshold used in identifying the significant events: FDR < 0.001, |IncLevel Difference| >= 0.15.

Regardless of which tissue was analyzed, the primary changes in alternative splicing relative to control tissue were increases in the number of skipped exons (SE) events (**Figure 4A**). FECD_REP or FECD_NR tissues showed more splicing changes than Pre_S tissue. The greater number of splicing changes is consistent with the extensive cellular degeneration observed in late stage disease. However, while not as many as in tissue from advanced disease, ∼450 changes in alternative splicing were observed in Pre_S tissue.

313 of the alternative splicing events in Pre_S tissue involved exon skipping (**Supplemental Table 3**). Heatmap evaluation of 313 skipped exons in Pre_S tissue revealed that the genes hosting the skipped exon events clustered most closely with FECD_REP tissue (**Figure 4BC**). 132 skipped exon events were shared between Pre_S and FECD_REP tissue, compared to only 28 were shared between Pre_S and FECD_NR tissue (**Figure 4B**), consistent with the hypothesis that the changes in Pre_S tissue foreshadow the alterations observed in FECD-REP.

While not nearly as frequent, other forms of alternative splicing events, alternative 5’ splice site (A5SS), alternative 3’-splice site (A3SS), mutually exclusive exon (MXE) and retained intron (RI), also showed that the Pre_S cohort was more similar to the FECD_REP group than the FECD_NR cohort (**Supplemental Figure 3A-D**). Taken together, these data demonstrate that splicing changes in Pre_S tissue are precursors to the changes in FECD_REP - but not FECD_NR - late stage disease.

We then separated the top 25 skipped exon events in Pre_S tissue, which are shared with FECD_REP for evaluation. As with the overall group of 313 events, the splicing of these genes was much more like FECD_REP tissue than to control or FECD_NR tissue (**Figure 4D**). Of the top 25 skipped exon events shared by FECD_REP and Pre_S tissue, approximately half are also seen in tibialis anterior muscle of myotonic dystrophy type 1 (DM1) subjects (32) (**Figure 4 DE, Supplemental Figure 3E-G**). This similarity, in spite of the comparison being made between samples from different tissues, suggests that the expanded CUG repeats in DM1 and FECD_REP have a common mechanism for producing splicing changes and that this mechanism is activated in Pre_S tissue.

Notable genes that show changes in splicing include the splicing factors *MBNL1* and *MBNL2*. Splicing of these genes was also changed in tissue from a myotonic dystrophy mouse model (32) and in a comparison of FECD_REP and FECD_NR tissue by Fautsch and colleagues (33, 34). Changes in *MBNL1* and *MBNL2* splicing in Pre_S (**Supplemental Table 3**) cells may trigger other splicing changes and eventually lead to the larger scale change that characterizes late stage disease.

### Changes in alternative splicing are quantitatively similar in Pre_S and FECD_REP tissue

We experimentally validated changes in splicing and levels of gene expression for Pre_S relative to control tissues (**Figure 5**). Genes were chosen for validation based on the significance of altered expression and the potential biological role of the gene expression suggested by pathway analysis (**Figure 8**).

**Figure 5.**
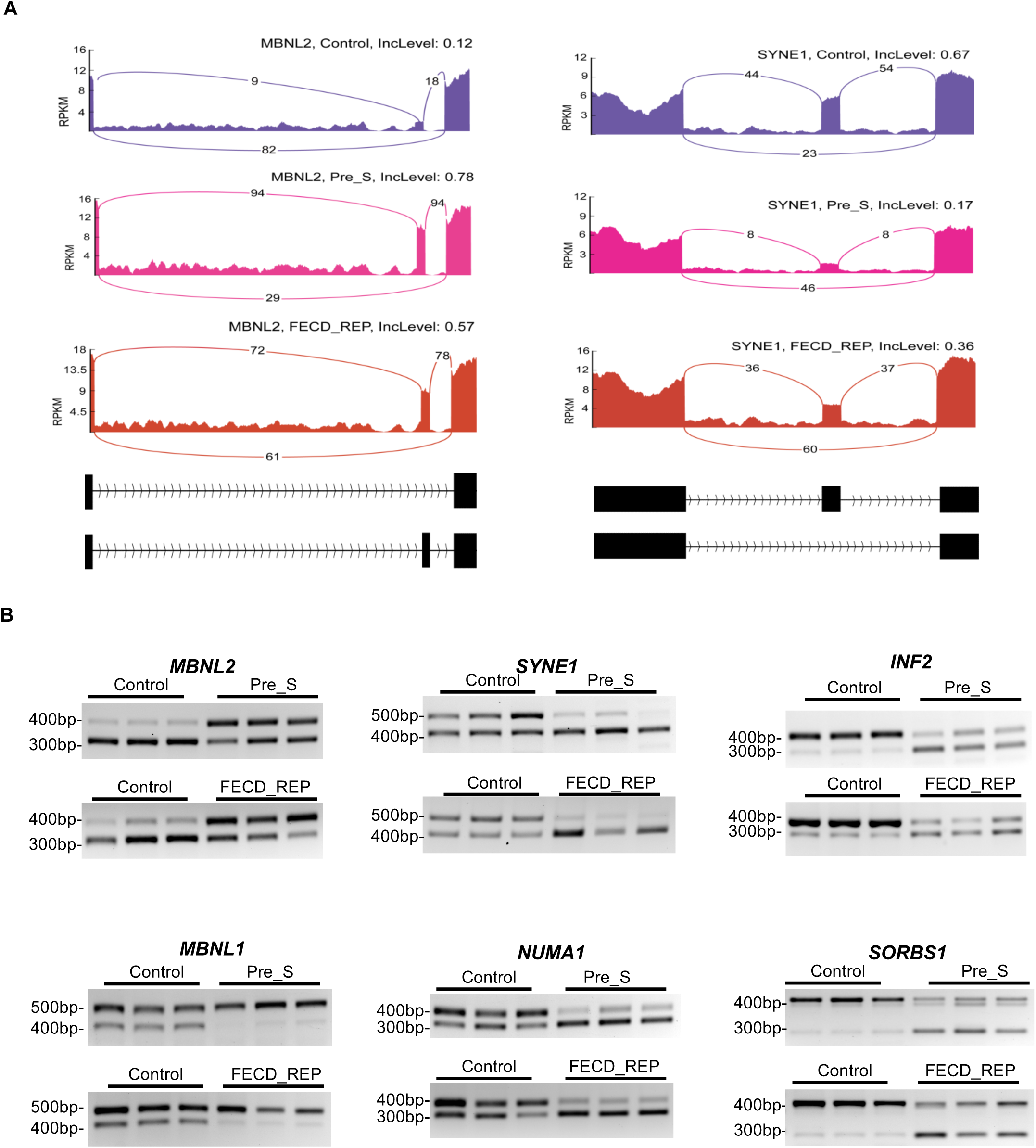
Differential RNA splicing is similar in Pre_S and FECD_REP tissue. **(A)** Representative Sashimi plots showing changes in alternative splicing for *MBNL2* and *SYNE1* transcripts in Control versus Pre_S and FECD_REP tissues. **(B)** Reverse-transcriptase PCR evaluation of RNA splicing changes in Control, Pre_S and FECD_REP tissue. All measurements used surgically prepared corneal endothelium from disease patients or donors detailed in Supplemental Table 1.

We used reverse-transcription PCR to evaluate changes of splicing for six genes, *INF2*, *NUMA1*, *SORBS1*, *SYNE1*, *MBNL1*, and *MBNL2* (**Figure 5**). INF2 protein associates with microtubules, and may affect cell shape. NUMA1 protein is component of the nuclear matrix and may affect mitotic spindle organization and proper cell division. *SORBS1* was chosen for its protein product’s involvement in cell adhesion and extracellular matrix. *SYNE1* encodes nesprin-1 which links the nuclear membrane to the actin cytoskeleton and may also affect cell morphology. *MBNL1* and *MBNL2* were chosen because potential alterations in their expression might feed back into even greater changes in splicing.

The changes in splicing predicted by global RNAseq analysis were confirmed by visual inspection of genes using Sashimi plots (Figure 5A) and validated by semi-quantitative analysis using reverse-transcriptase PCR (RT-PCR) (Figure 5B) using monolayer human corneal endothelial tissue. Pre_S tissue showed the changes in splicing predicted by our RNAseq data (**Figure 5**). Both Sashimi plots and semi-quantitative reverse transcriptase PCR (RT-PCR) analysis reveal that the absolute magnitude of splicing changes are similar in FECD_REP late stage disease tissues. The amount of change in splicing in Pre_S tissue is substantial, similar to that observed in late stage tissue. This observation suggests that missplicing of key genes does not gradually increase as disease symptoms progress, rather substantial missplicing is a leading indicator of disease.

### Differential gene expression

We compared gene expression level changes in Pre_S, FECD_REP, and FECD_NR tissue cohorts relative to control tissue. To classify changes as significant, we used the adjusted *p*-value generated by Cuffdiff, one of the programs within Cufflinks suite, as the determining metric. Changes with a *p*-value less than 0.05 were deemed significant and included in our analysis.

All sample cohorts showed expression changes relative to control tissue (Pre_S: 215, FECD_REP: 1330; FECD_NR: 696) (**Figure 6A and 6B**). The greater number of gene expression changes in the FECD_REP and FECD_NR tissues is consistent with the severe cellular phenotype observed in late stage disease. 602 out of the 696 genes differentially expressed in the FECD_NR tissues were also found in the FECD_REP group suggesting significant overlap of the common final molecular genetic mechanisms in the two forms of late-stage disease.

**Figure 6.**
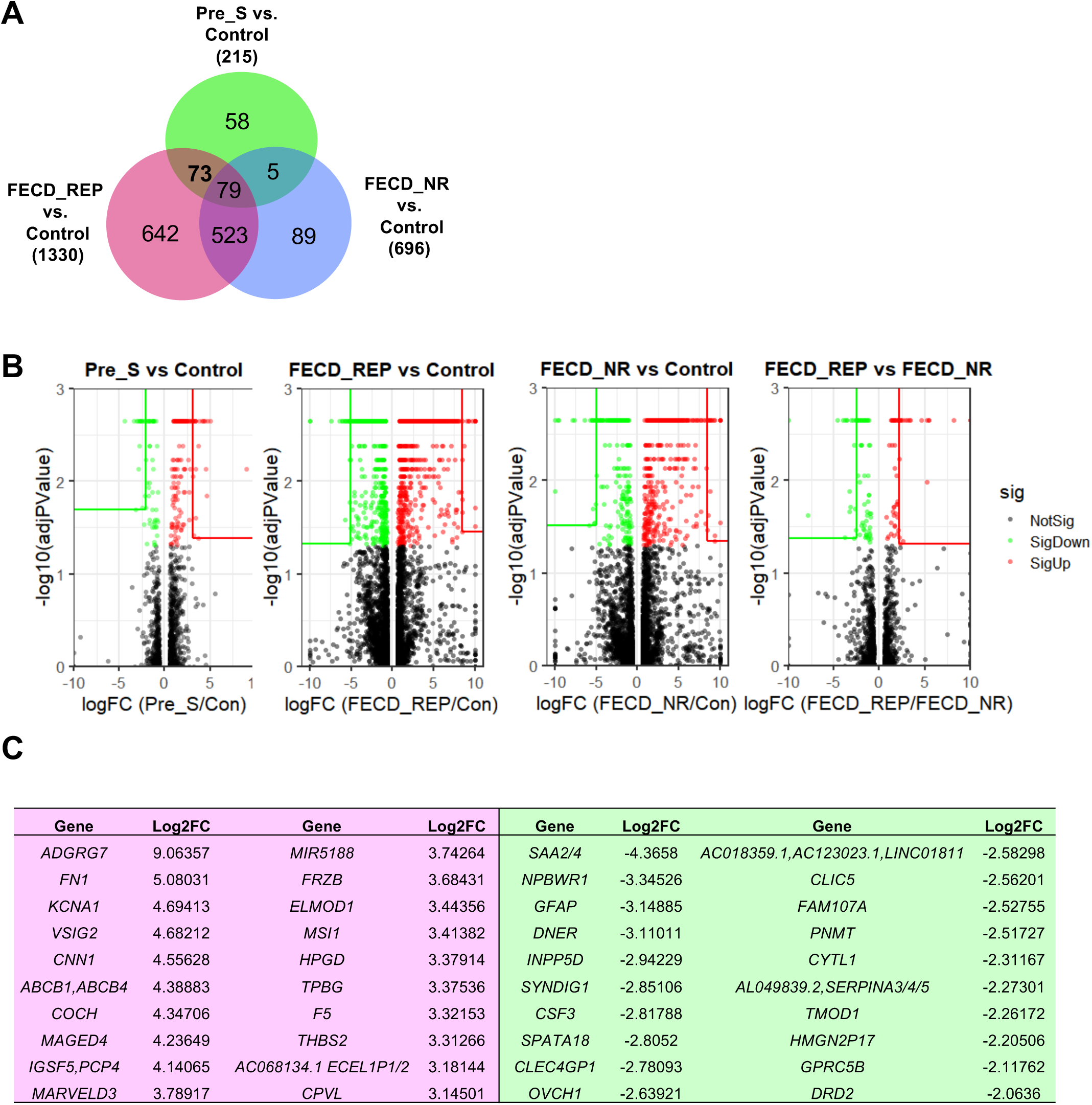
RNAseq analysis of gene expression. (**A**) Overlap for gene expression of Pre-S, FECD_REP, and FECD_NR cohorts. Numbers indicate the number of differentially expressed genes identified in each individual region. (**B**) Volcano plots for Pre_S vs. Control, FECD_Rep vs. Control, FECD_NR vs. Control and FECD_REP vs. FECD_NR and top genes in each comparison. (**C**) Top 20 upregulated and downregulated genes in Pre_S vs. control comparison.

Pre_S tissue had 215 genes with significantly altered expression levels relative to control tissue. Only five changes in gene expression were uniquely shared between Pre_S and FECD_NR tissue, compared to 73 shared changes with FECD_REP. (**Figure 6A**). The closer relationship between Pre-S and FECD_REP is consistent with our splicing data (**Figures 4 and 5**) and supports the conclusion that patterns of gene expression in mutant expanded repeat cells are established long before symptoms or disease findings are observable.

Volcano plots allow a global overview of individual gene expression changes. They are useful for visualizing patterns of changes and identifying “outlier” genes that combine highly significant changes in gene expression with higher fold changes. The fold change among top genes are less in Pre_S than in FECD_REP or FECD_NR and the identity of top genes differ (**Figure 6B and 6C, Supplemental Tables 4-6, Supplemental Figure 4**). This finding is consistent with the severity of late stage disease and the disruption of many gene expression programs.

Evaluation of Pre_S tissue may provide a window to identify early gene drivers before symptom-driven secondary changes in gene expression overwhelm analysis. The top twenty differentially over-expressed genes identified in Pre_S tissue (**Figure 6C**) include genes involved in the extracellular matrix and its assembly, cochlin (*COCH*) (35, 36), fibronectin (*FN1*) (37), and thrombospondin (*THBS2*) (38).

### Quantitative analysis of gene expression changes

We used quantitative PCR (qPCR) to confirm the changes in the level of gene expression detected by RNAseq in both Pre_S and FECD tissue (**Figure 7**). qPCR measurements of corneal endothelial tissues were challenging because of the limited amount of material available, but we could compare expression of eight genes identified in our RNAseq data, *FN1*, *COL4A2*, *COCH*, *CTGF*, *MSI1*, *LUM*, *KDR*, and *SOD3*. Four of these genes, *FN1*, *COL4A2*, *CTGF*, *KDR*, were within the fibrosis pathway. *COCH* and *LUM* encode extracellular matrix proteins, *MSI1* encodes a RNA binding protein/splicing factor, and SOD3 protein is related to oxidative stress. The observed changes in gene expression confirm our RNAseq results (**Figure 6B and Supplemental Table 7**).

**Figure 7.**
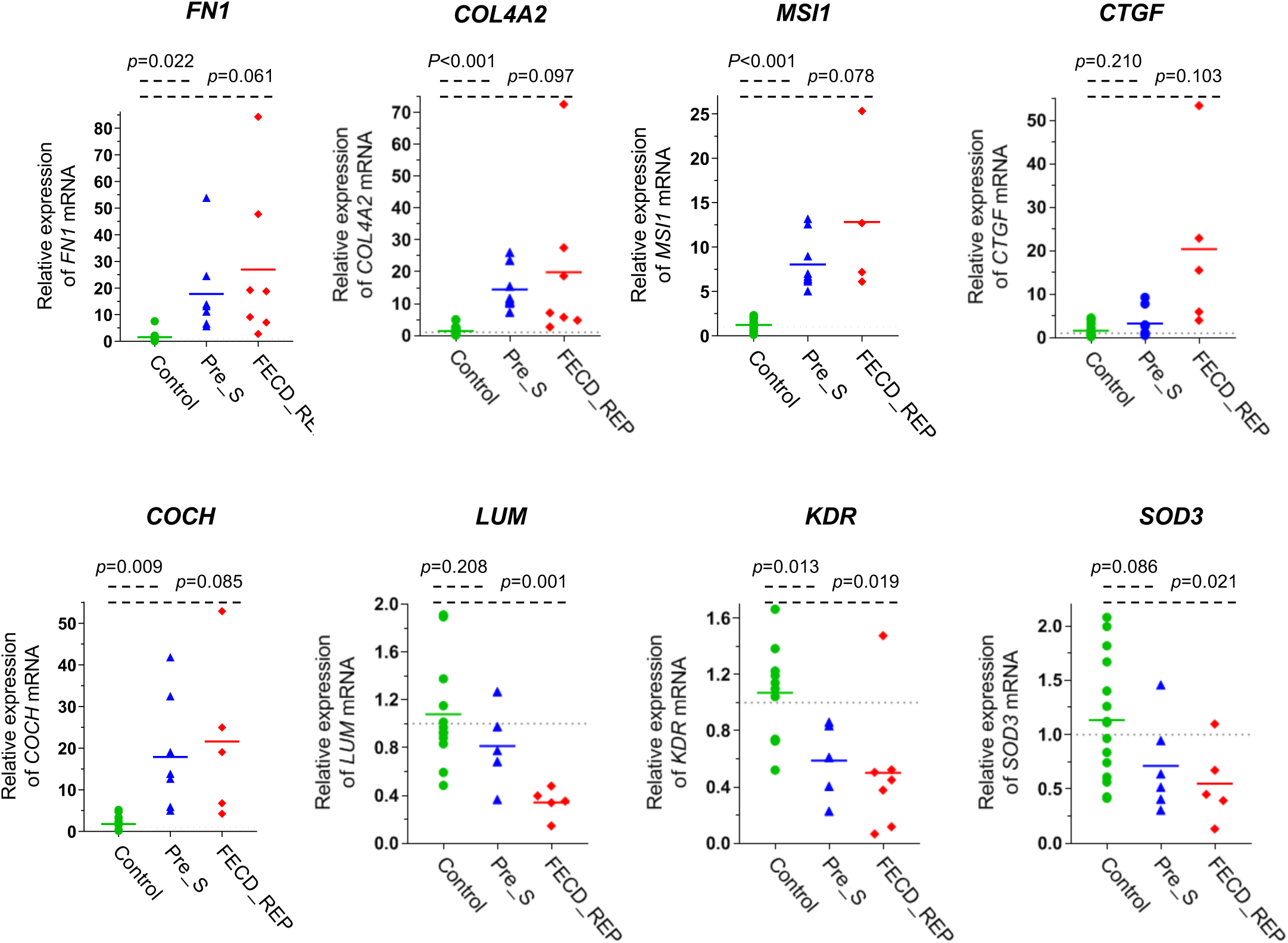
Changes in gene expression in Pre_S and FECD_REP tissue. qPCR evaluation of gene expression levels in control, Pre_S, and FECD_REP tissue. All measurements used surgically prepared single cell monolayer corneal endothelium from disease patients or donors detailed in Supplemental Table 1.

### Pathway analysis

To elucidate the potential impact of changes in the expression of individual genes on physiologic processes, we applied Ingenuity Pathway Analysis (IPA) to our RNAseq data. Overwhelmingly, the top common canonical pathway was hepatic fibrosis/hepatic stellate cell activation (39) (**Figure 8A)**. Involvement of the hepatic fibrosis pathway genes in FECD_REP and FECD_NR is consistent with the observed accumulation of extracellular matrix (ECM) in advanced FECD with thickening of Descemet’s membrane with focal excrescences (guttae) and with previously reported RNA measurements using tissue from late stage disease (40). Our FECD transcriptome data indicates robust ECM production in late-stage disease possibly regulated by transforming growth factor-β (TGF-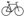), the most potent fibrogenic cytokine released by a number of cell populations in the body including the liver (39) (**Supplemental Figure 5AB**).

Activation of the fibrosis pathway was also observed in Pre_S tissue (**Figure 8AB, Figure 9**). Genes that showed statistically significant increases in expression include fibronectin *FN1*, one of the highest differentially expressed genes in Pre_S tissue (**Figure 7**). Other genes include connective tissue growth factor (*CTGF*) and four members of the collagen alpha chain family including *COL1A2* which is also abundant in liver fibrosis (**Supplemental Table 7**) (41). Kinase insert domain receptor (KDR, also known as vascular endothelial growth factor receptor-2) showed decreased expression. Relevant in the fibrosis pathway, KDR protein interacts with VEGF to mediate vascular endothelial cell proliferation. These genes are also disturbed in advanced disease - another indication that gene expression programs associated with fibrosis are activated in presymptomatic carriers prior to observable symptoms of disease.

**Figure 8.**
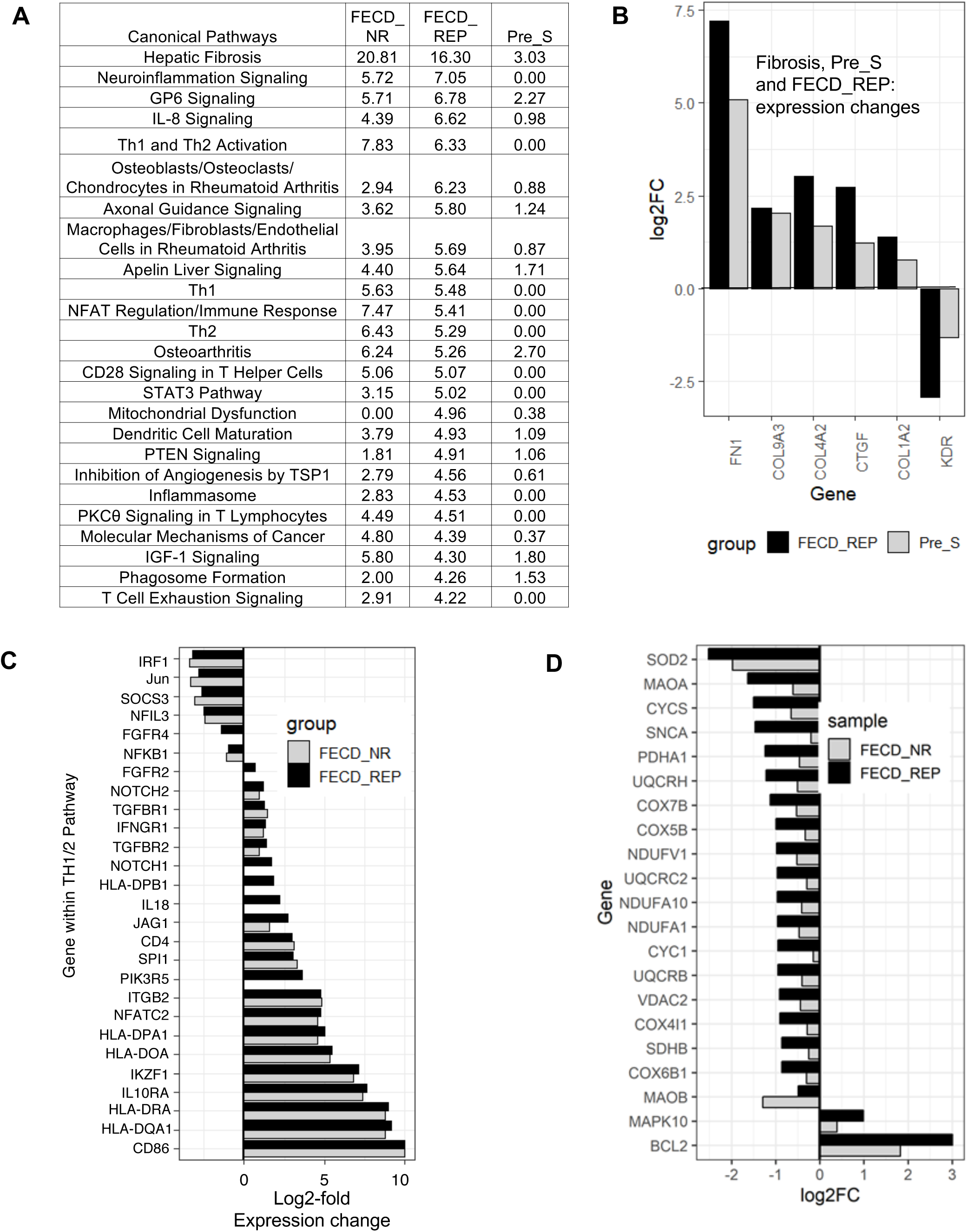
Pathway analysis. (**A**) Top 25 pathways based on FECD_REP. The *p*-values for three cohorts for each pathway are shown in the last three columns. (**B**) Shared differentially expressed genes in Fibrosis pathway between Pre-S and FECD_REP. (**C**) Changes in Th1/2 pathway genes for FECD_NR and FECD_REP. (**D**) Significant expression level changes in genes involved in mitochondrial dysfunction seen in FECD_REP but not in FECD_NR. FPKM > 1.5, fold change > 1.5, FDR <=0.05.

**Figure 9.**
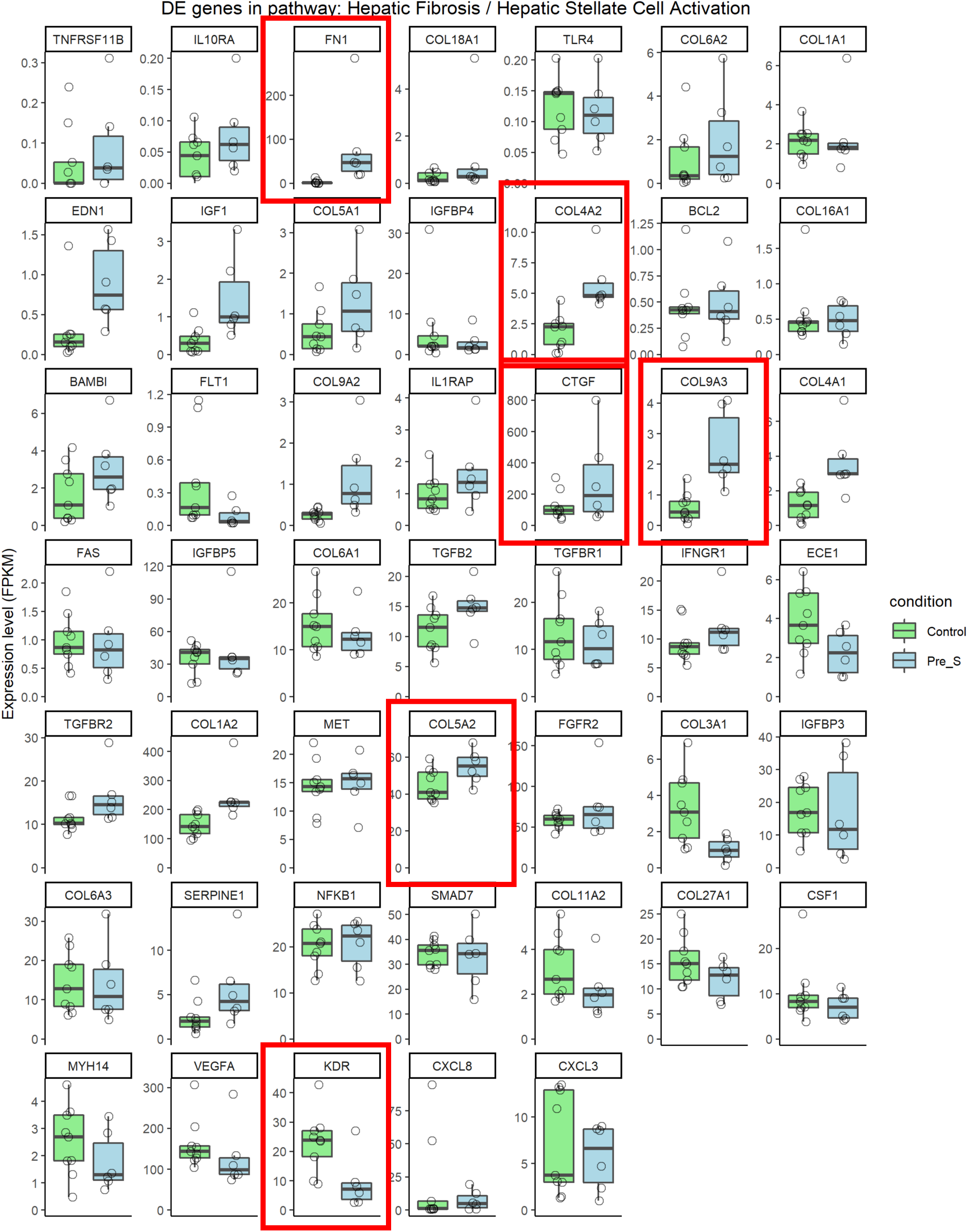
Expression profile of genes in Hepatic fibrosis pathway for Pre_S cohort. FPKM > 1.5, fold change > 1.5, FDR <=0.05. The six largest fold changes in gene expression (Figure 8) are outlined in red.

The pathways underlying FECD_REP and FECD_NR advanced stage disease had significant overlap. There were numerous shared pathways implicating the immune system related to helper T cell activation, signaling, and neuroinflammation. (**Figure 8A and 8C**). Marked overexpression of genes encoding proteins on the surface of antigen-presenting cells including the B7 protein, *CD86* (> 2,000 fold increase), and class II major histocompatibility proteins (> 250 fold) both required for these cells to activate helper T cells implicate the immune system in both FECD groups in late-stage disease (**42**) (**Figure 8C, Supplemental Figure 5C-E, Supplemental Table 4 and 5**).

A few molecular pathways were changed in FECD_REP but not FECD_NR (**Figure 8D**). The canonical pathway related to mitochondrial dysfunction showed the large difference with *p* values of 10^-6^ and zero respectively for FECD_REP and FECD_NR (**Figure 8A, Supplemental Figure 5FG**). Decreased expression of oxidative phosphorylation genes was more pronounced in FECD_REP compared to the FECD_NR (**Figure 8D, Supplemental Figure 5H)**.

## Discussion

Identifying the early drivers of late-onset disease is important for understanding disease progression and developing therapeutics. Studying early drivers, however, is often not practical because pre-symptomatic tissue is difficult to obtain. Because the expanded CUG repeat mutation within *TCF4* intron 2 that causes FECD_REP is so prevalent (3% of the Caucasian population), significant numbers of pre-symptomatic samples can be obtained from individual donors positive for the CUG expansion. These tissues, together with FECD_REP, FECD_NR, and control tissues (**Figure 1 and 2**) provide an advantageous model for better investigating the early links between expanded trinucleotide repeat mutations and disease.

The goal of this study was to understand whether the expanded CUG repeat was changing gene expression in Pre_S tissue and how such changes might relate to the gene expression and phenotypic changes known to occur in late-stage FECD.

### FECD is a disease of mutant RNA

FECD_REP is caused by an expanded CUG trinucleotide repeat within mutant *TCF4* intronic RNA (**Figure 10**) (15–18). Remarkably, FECD is also caused by the mutant CUG expanded repeat within the 3’-untranslated region of the *DMPK* gene is also associated with myotonic dystrophy (19–21). Unlike other trinucleotide repeat diseases where the contribution of mutant RNA is debated, data demonstrating that CUG RNAs expressed from two different genes are both responsible for FECD offers strong support for RNA playing a central role in the molecular origins of the disease.

**Figure 10.**
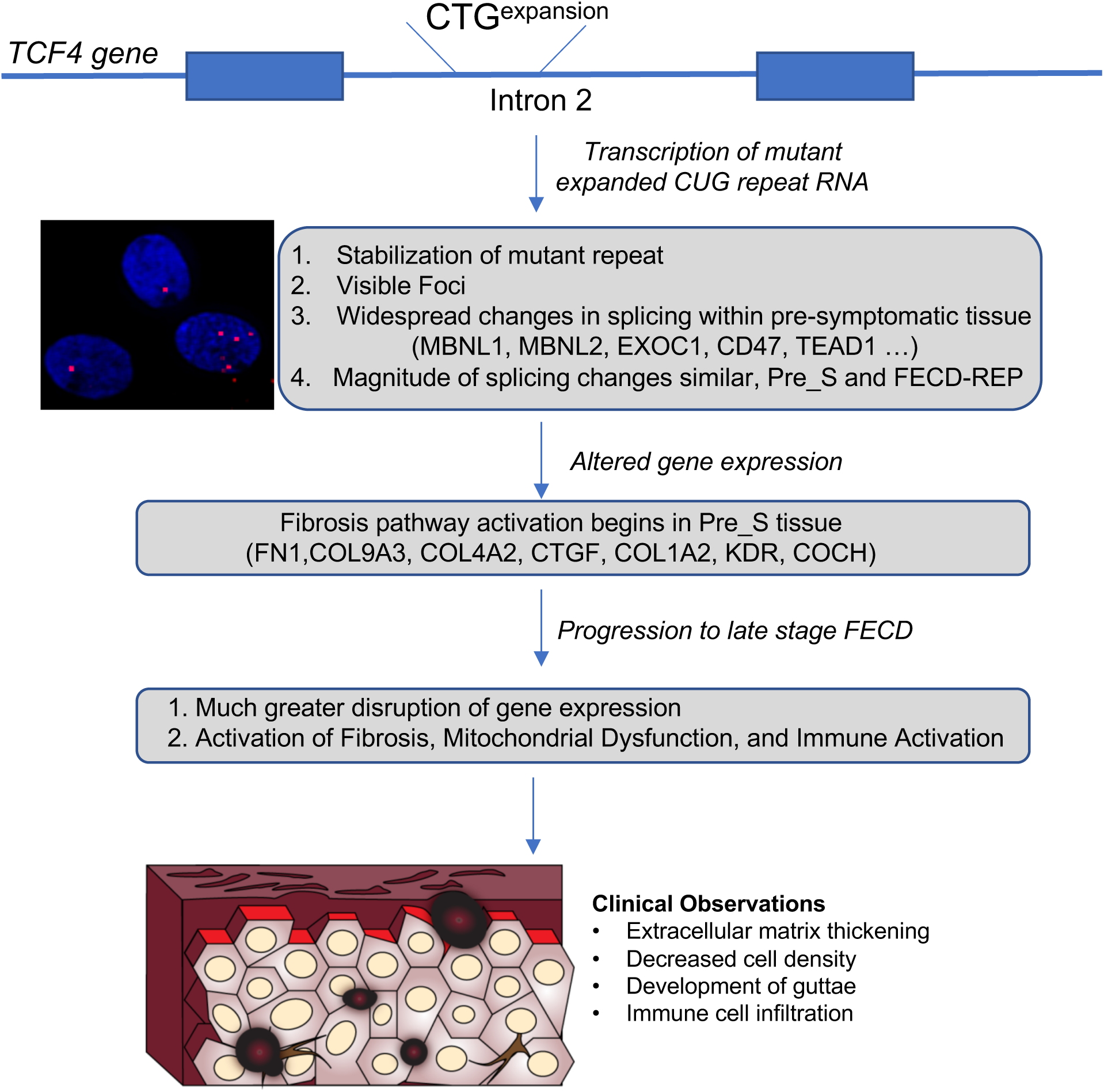
Schematic diagram of FECD molecular and disease progression from CTG expanded repeat mutation at the *TCF4* locus to advanced FECD. Initially the CTG expansion expresses the CUG repeat RNA. Because the expanded repeat mutation that causes FECD_REP is a relatively common genetic mutation, Pre_S tissue is available for analysis. Pre_S tissue appears normal upon clinical observation. However, in Pre_S tissue foci can be detected, we observe changes in splicing and gene expression, the mutant intronic repeat is stabilized, and early signs of fibrosis pathway activation are apparent. In late stage disease, more pronounced changes in splicing and gene expression accompany clinically observable systems and loss of vision.

The expanded CUG can be detected by fluorescent in situ hybridization (FISH) as RNA foci. These RNA foci are a hallmark of both Pre_S and FECD_REP corneal endothelium (**Figure 10**) (43). While Pre_S and FECD_REP tissue both possess the RNA trigger for FECD, Pre_S tissue is visually indistinguishable from control tissue upon specular imaging. By contrast, FECD_REP tissue is dramatically different from control tissue, with reduced cell density and the formation of focal collagen accumulations known as guttae.

The FECD CUG repeat RNA is present at only a few (<10) copies per cell in disease tissue (44). It is likely that each “foci” detected by FISH is a single RNA molecule. A low copy number for a disease-causing RNA has also been observed for myotonic dystrophy type 1(DM1) (45) and *C9orf72* ALS/FTD (46). We find that the CUG repeat expansion stabilizes *TCF4* intronic RNA in a corneal cell-specific manner (**Figure 3**). This enhanced stability may contribute to an ability of a small number of RNA molecules to bind protein sufficiently to affect overall function in cells and eventually produce observable symptoms that characterize a delayed onset disease like FECD.

Previous work has suggested that the CUG repeat within intronic *TCF4* and other microsatellite expansions are associated with intron retention (47). Our detection of nuclear CUG RNA foci by FISH and upstream intronic RNA due to its increased half-life may be compatible with a disease model where the *TCF4* intron with the expanded repeat is spliced out, forms a linear stable intronic sequence RNA, and undergoes preferential 3’ to 5’ exonuclease degradation in the nucleus.

There is no definitive mechanistic insight into how the relatively rare expanded CUG repeat RNA can cause FECD. How one or a few copies of RNA triggers widespread changes in gene expression and late onset disease remains a major unanswered question. However, studies of myotonic dystrophy have suggested that the CUG repeat binds members of the muscleblind-like (MBNL) protein family. MBNL proteins are splicing factors and their sequestration reduces the concentration of free MBNL in cell nuclei and affects splicing. Overexpression of MBNL1 can reverse RNA missplicing and myotonia in a DM1 mouse model (31). MBNL is also associated with mutant CUG RNA in FECD cells and tissue (28–30). Blocking the CUG repeat region using antisense oligonucleotides can reverse missplicing in DM1 (caused by a CUG repeat within the *DMPK* gene) (48–51) and FECD (44, 52) tissues. These reports support the hypothesis that the mutant RNA may act by binding to MBNL and affecting splicing (**Figure 10**).

We have recently reported that relatively little MBNL protein is in the nuclei of FECD_REP cells and human tissue (53). Using quantitative protein titrations against a known standard, we calculated that there were 65,000 copies of MBNL1 and MBNL2 per cell and less than 2000 copies were present in cell nuclei. Low copy numbers for MNBL in the nuclei of affected tissue are consistent with the hypothesis that even a small amount of mutant expanded CUG repeat RNA may be sufficient to affect the available pool of MNBL protein. A reduction in available MBNL protein would produce the alterations of splicing that are a hallmark of FECD_REP disease (**Figures 4 and 5**). It is also possible that the MNBL:mutant RNA interaction may nucleate additional protein or RNA interactions to amplify disruptive effects on gene expression and changes in alternative splicing.

### Expanded CUG mutant RNA causes splicing changes in presymptomatic tissue

Many of the alterations in splicing observed in FECD_REP tissue also define gene expression in presymptomatic tissue, Pre_S (**Figures 4 and 5**). The splicing factors MBNL1 and MBNL2 are among the genes showing altered splicing in Pre_S samples and these changes may amplify other splicing changes to push corneal endothelial cells towards full blown FECD.

The similarity of alternative splicing changes between symptomatic FECD_REP and presymptomatic Pre_S tissue is much greater than that between Pre_S and FECD_NR samples. The data suggest that there are fundamental differences in the origins of the two forms of FECD. While their origins differ, late-stage FECD_REP and FECD_NR converge at a common set of clinical findings.

The data also suggest that splicing changes and perturbation of extracellular matix (ECM) genes seen in FECD_REP late stage disease tissue begin to be observed long before symptoms are observed during standard clinical evaluation. The molecular trigger, mutant RNA, and early molecular changes, altered splicing and observable RNA foci, co-exist in cells that appear to have a normal phenotype. The changes in the magnitude of RNA splicing between control tissue and either Pre_S or late stage FECD_REP samples are similar. This observation suggests that the mutant RNA triggers the splicing changes in key genes independent of the progression of disease.

The finding that splicing is an early trigger has important implications for the development of agents to treat FECD. It is reasonable to expect that such agents would be most effective when administered early in disease progression prior to exuberant production of ECM with degeneration of the corneal endothelium and activation of the immune system. During drug development it should be possible to monitor the changes in alternative splicing and expression of ECM biomarkers caused by expression of the mutant trinucleotide repeat and rank drug candidates by their ability to return splicing to a more normal state – agents that reverse the splicing defect would be promising development candidates. Monitoring splicing of key genes would offer a rapid and definitive assay for screening compounds. We have previously shown that synthetic oligonucleotides complementary to the CUG repeat can reverse the splicing defect (43).

Recently, Fautsch and colleagues reported that some individuals without previously noted guttae who possessed the expanded repeat but developed “non-FECD corneal edema” do not show changes in alternative splicing (33). These results might appear to be in conflict with our observation that all CUG expanded repeat individuals exhibited substantial changes in alternative splicing, including many shared between Pre_S and FECD_REP individuals (**Figure 4**). We note that the expanded repeat positive individuals with corneal decompensation without findings of FECD were between 67 and 83 years old, much older than any individual in our Pre_S cohort. As postulated by Fautsch and colleagues, it is possible that these individuals possessed protective mutations that might prevent splicing changes and block the molecular events leading to symptomatic FECD.

### The fibrosis pathway is activated in Pre_S tissue

In the clinic, late stage FECD is a disease of cellular degeneration and aberrant extracellular matrix deposition (**Figure 10**). Previous studies of FECD_REP tissue have supported activation of the fibrosis pathway as a primary cause for late stage disease pathology (30). We confirmed fibrosis as the highest ranked canonical pathway in both late stage FECD_REP and FECD_NR (**Figure 8A**). The fibrosis pathway is also activated in Pre_S tissue (**Figure 8A,B, Supplemental Table 7**). These results demonstrate that changes in splicing, changes in gene expression, and changes in a key disease pathway begin years before symptoms are observed.

Among the top ten genes in pre-symptomatic tissue versus control tissue are cochlin (*COCH*) and fibronectin (*FN1*) with >16 and >32 fold change increases respectively. Cochlin is a secretory extracellular matrix protein originally identified in the cochlear cells of the inner ear (35). *COCH* has also been found to expressed by the endothelial cells the trabecular meshwork of subjects with primary open-angle glaucoma (POAG), another common age-related degenerative disorder (36). There appears to be a convergence of two age-related disorders of the anterior segment of the eye mediated by *COCH*. Primary open-angle glaucoma may be more prevalent in patients with advanced FECD (54). Primary open angle glaucoma is also a disease mediated by transforming growth factor-β which increases aqueous humor outflow resistance by dysregulation of ECM genes in the endothelial cells lining the trabecular meshwork (55). One area for future research will be to determine whether *COCH* or other ECM genes with altered expression in Pre_S tissue can serve as biomarkers for early detection of FECD or to facilitate the monitoring of clinical trials.

### Late stage FECD tissue is characterized by changes in immune cell-related and mitochondrial dysfunction pathways

Both FECD_REP and FECD_NR tissues show activation of genes related to immune system required for helper T cell activation, signaling, and neuroinflammation (**Figure 8AC, Supplemental Figure 5C-E and Figure 10**). In both groups, we detected a 2000-fold increase in *CD86* and marked upregulation of major histocompatibility genes (MHC) required by antigen presenting cells to activate helper T cells. This gene expression data along with the recent observation of cells with a dendritic morphology and positive for the hematopoietic marker CD45 in the endothelial tissue keratoplasty specimens of patients (39) suggest an important role for antigen presenting cells in late-stage disease in both forms of FECD.

The mitochondrial dysfunction pathway is activated in FECD_REP tissue with the expression of over twenty genes changed, little change was seen in FECD_NR tissue (**Figure 8D, Supplemental Figure 5F-H**). Further research will be needed to determine whether this difference plays a role in disease progression or response to treatment.

### Conclusion

FECD has many advantages as a model for understanding the origins of trinucleotide repeat disease because pre-symptomatic tissue is relatively accessible. Examination of Pre_S tissue reveals changes in gene expression that preview the more extensive changes in late stage disease. In particular, there is early activation of key genes associated with the fibrosis pathway, the pathway that defines the primary phenotype observe during advanced disease. Splicing patterns and levels of expression for key genes change decades prior to observation of the clinical manifestations of FECD_REP. Surprisingly, many changes in alternative splicing are similar in magnitude in Pre_S and advanced stage FECD_REP tissue. Many altered alternative splicing changes are shared with myotonic dystrophy, another disease caused by expanded CUG trinucleotide repeats, and it is possible that our findings will also be applicable to the genesis and temporal progression of other trinucleotide repeat diseases.

## Author Contributions

YC and JH planned experiments and analyzed data. HL, MK, and CX analyzed data. WB and CB provided surgical specimens. XG performed genotyping experiments. DRC and VVM initiated the project, supervised the project, and prepared the manuscript.

## Acknowledgements

The authors thank the patients for their participation in this study. We appreciate the support of Dwight Cavanagh and Donna Drurry at Transplant Services at UT Southwestern. We thank Albert Jun for generously sharing F35T cell line. We thank Chelsea Burroughs and Katherine Corey for helping with Figure 10 and Bethany Janowski, Keith Gagnon, Jay Horton, and Jerry Niederkorn for their helpful comments. This study was supported by grants R01EY022161 (VVM), P30EY030413 (VVM), and R35GM118103 (DRC) from the National Institutes of Health, Bethesda, MD, an unrestricted grant from Research to Prevent Blindness, New York (VVM), Harrington Scholar-Innovator Award from Harrington Discovery Institute (VVM), the Alfred and Kathy Gilman Special Opportunities in Pharmacology Fund (DRC), and the Robert A. Welch Foundation I-1244 (DRC). VVM is the Paul T. Stoffel / Centex Professor in Clinical Care. DRC is the Rusty Kelley Professor of Biomedical Science.

---

**Supplemental Figure 1.**
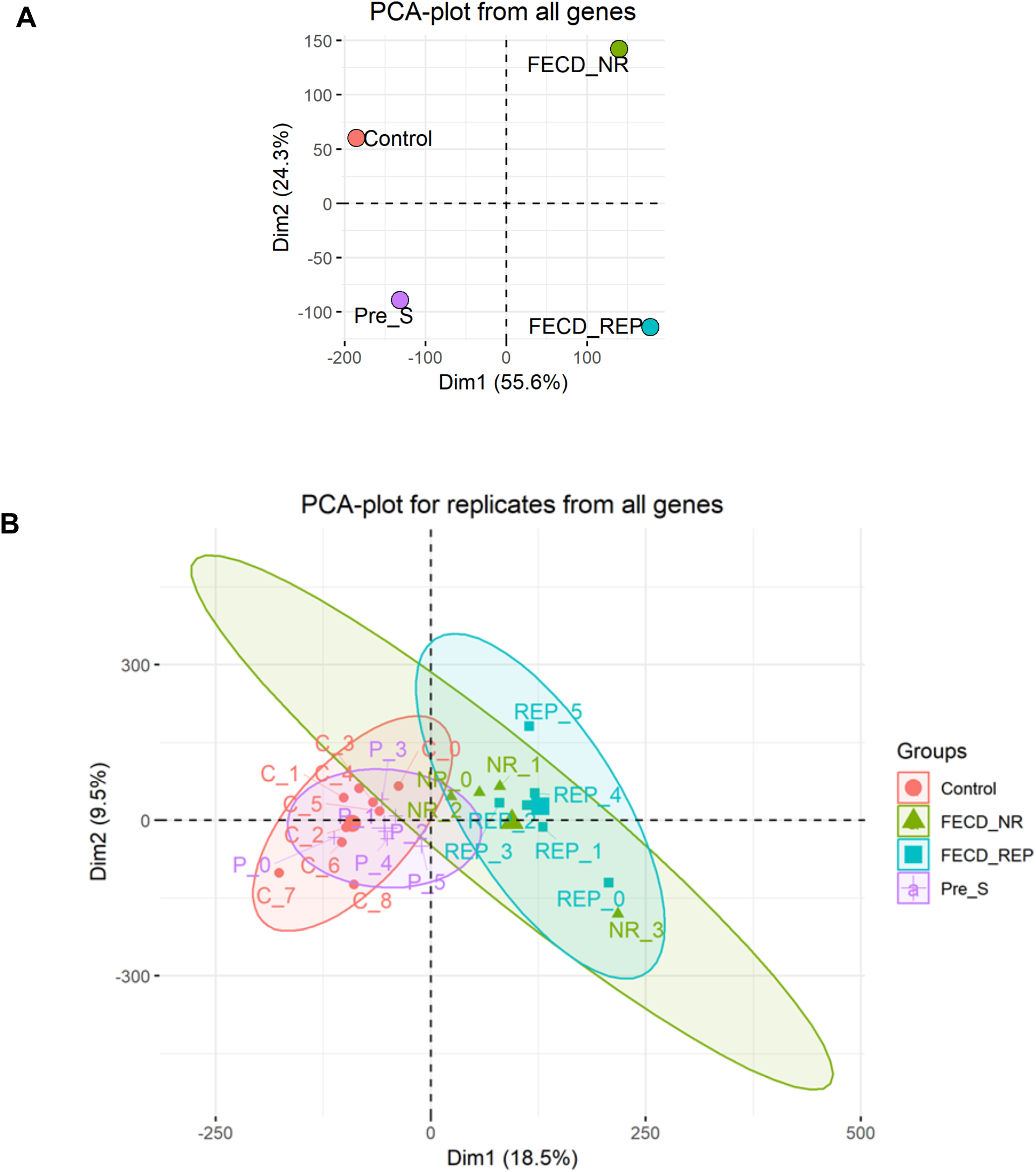
**Sample clustering**. (**A**) Principle Component Analysis (PCA) plot showing the group relationship based on gene expression for four groups of samples. **(B**) PCA plot for all the replicates.

**Supplemental Figure 2.**
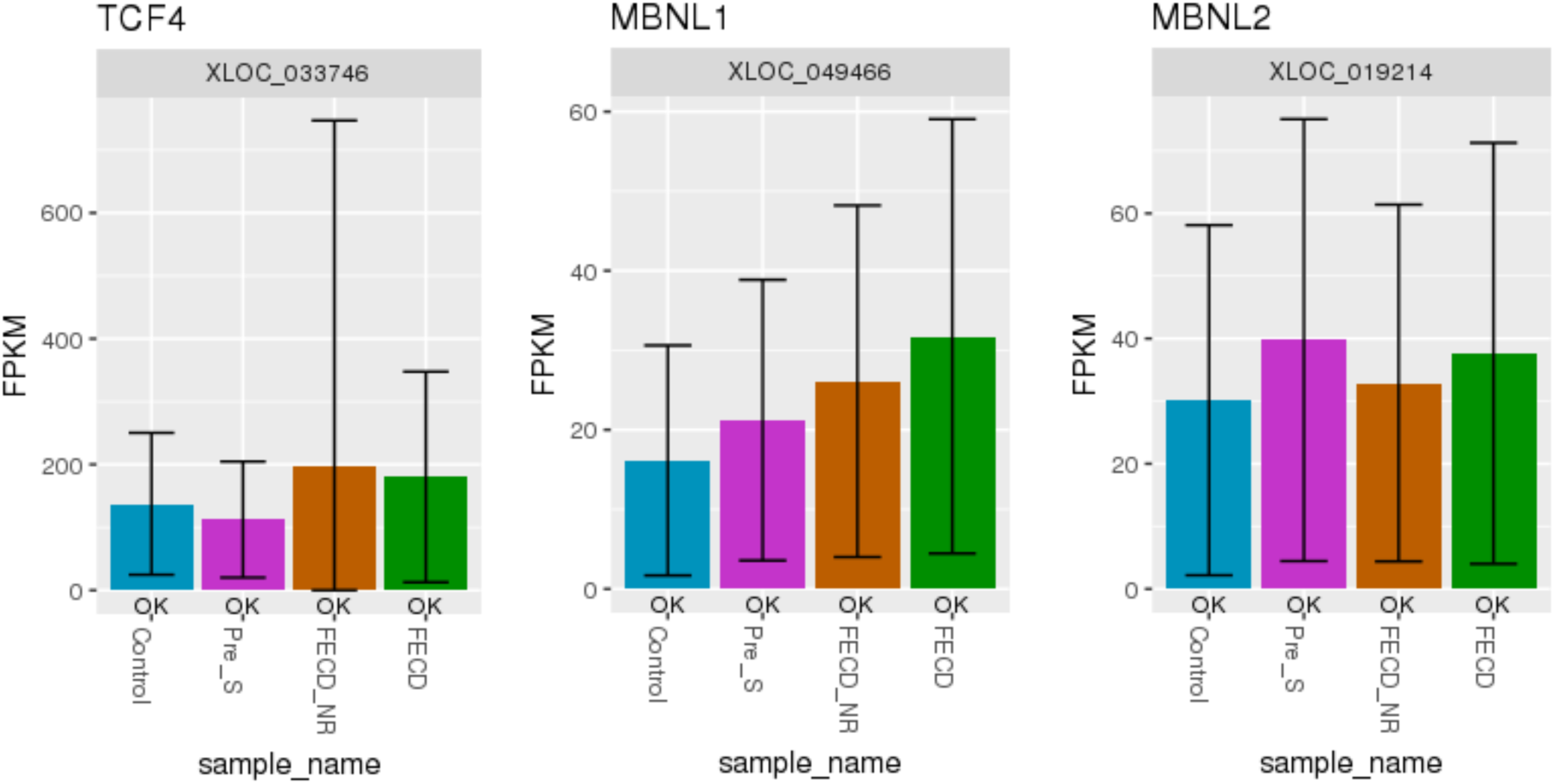
**Expression levels of *TCF4*, *MBNL1*, and *MBNL2* mRNA.** Measurements were made using RNAseq and corneal endothelial tissue samples from the four sample cohorts. No statistically significant differences were observed. Error bar stands for 95% confidence interval.

**Supplemental Figure 3.**
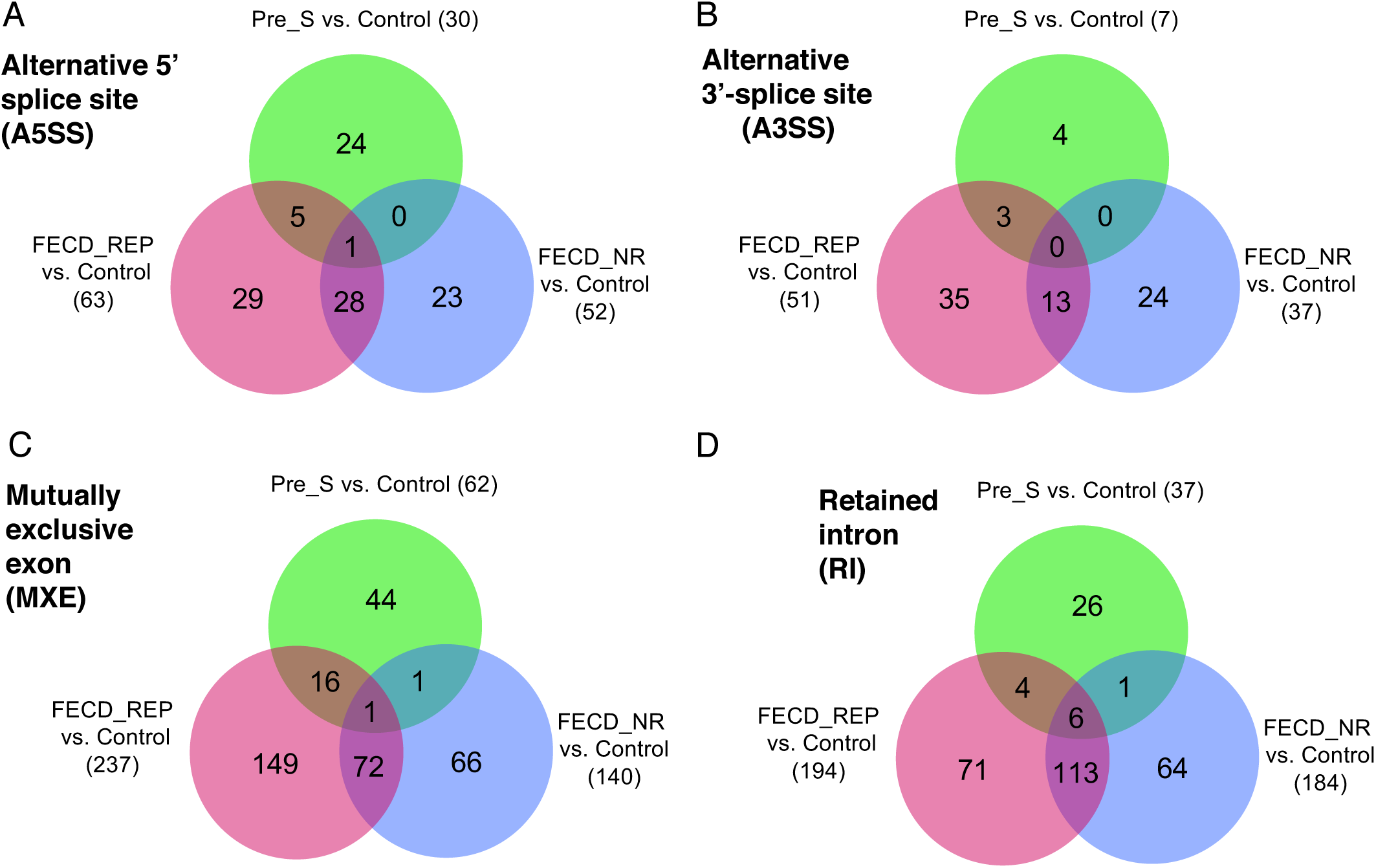
**RNAseq data demonstrates changes in splicing in pre-symptomatic, FECD_REP, and FECD_NR tissue.** (**A-D**) Alternative splicing events, A5SS, A3SS, MXE, RI in Pre_S, FECD_REP, and FECD_NR cohorts (relative to control). The threshold used in identifying the significant events: FDR < 0.001, |IncLevel Difference| >= 0.15.

**Supplemental Figure 3.**
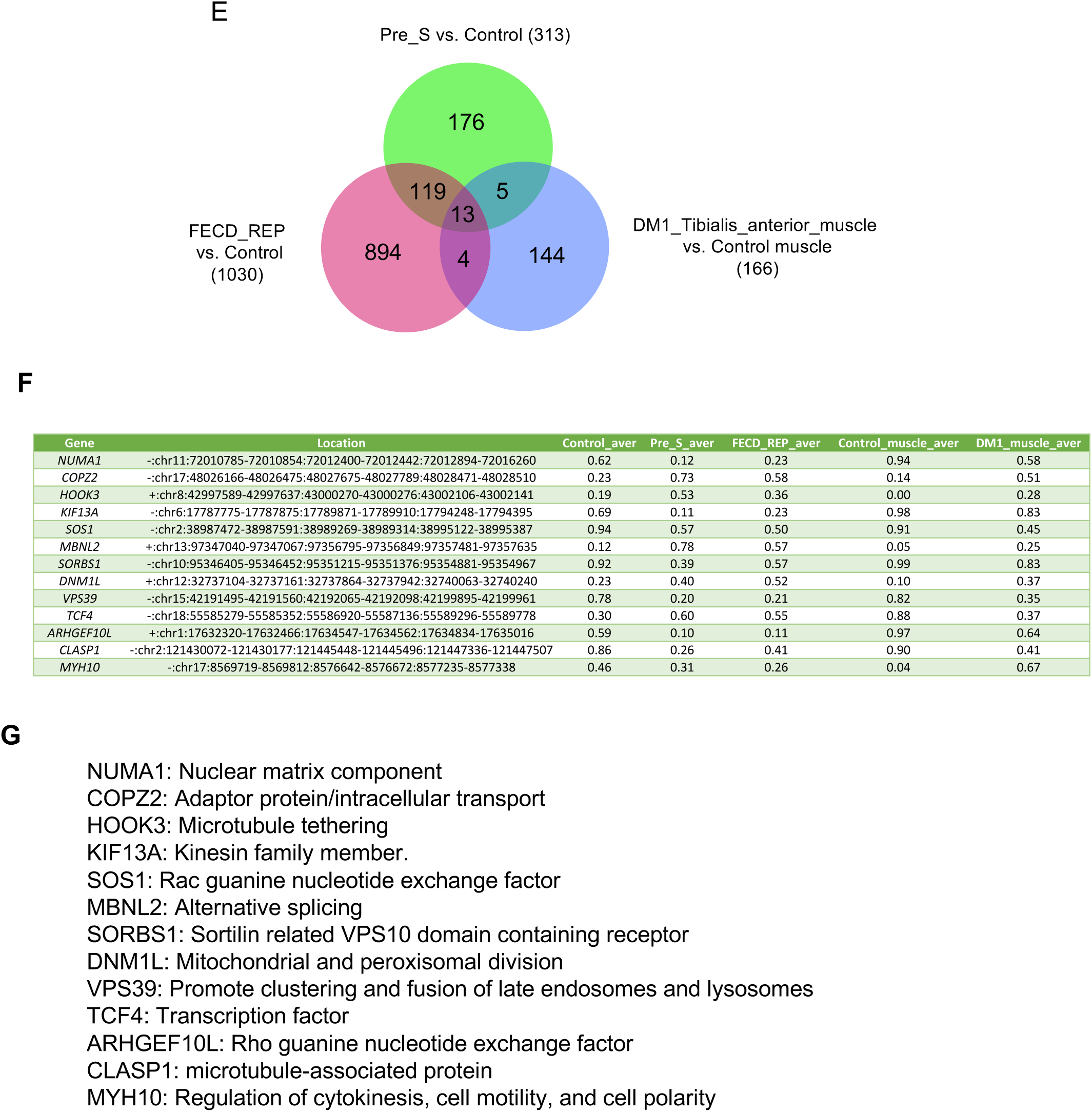
(**E**) Skipped exon events are shared among Pre_S, FECD_REP and DM1. (**F**) Details of 13 shared SE events among three groups, FECD_REP, Pre_S, and DM1. DM1 raw data was obtained from Gene Expression Omnibus (GSE86356) and analyzed similarly as the FECD data (visit DMseq.org for more information). (**G**) Basic functions of commonly shared genes listed in (F).

**Supplemental Figure 4.**
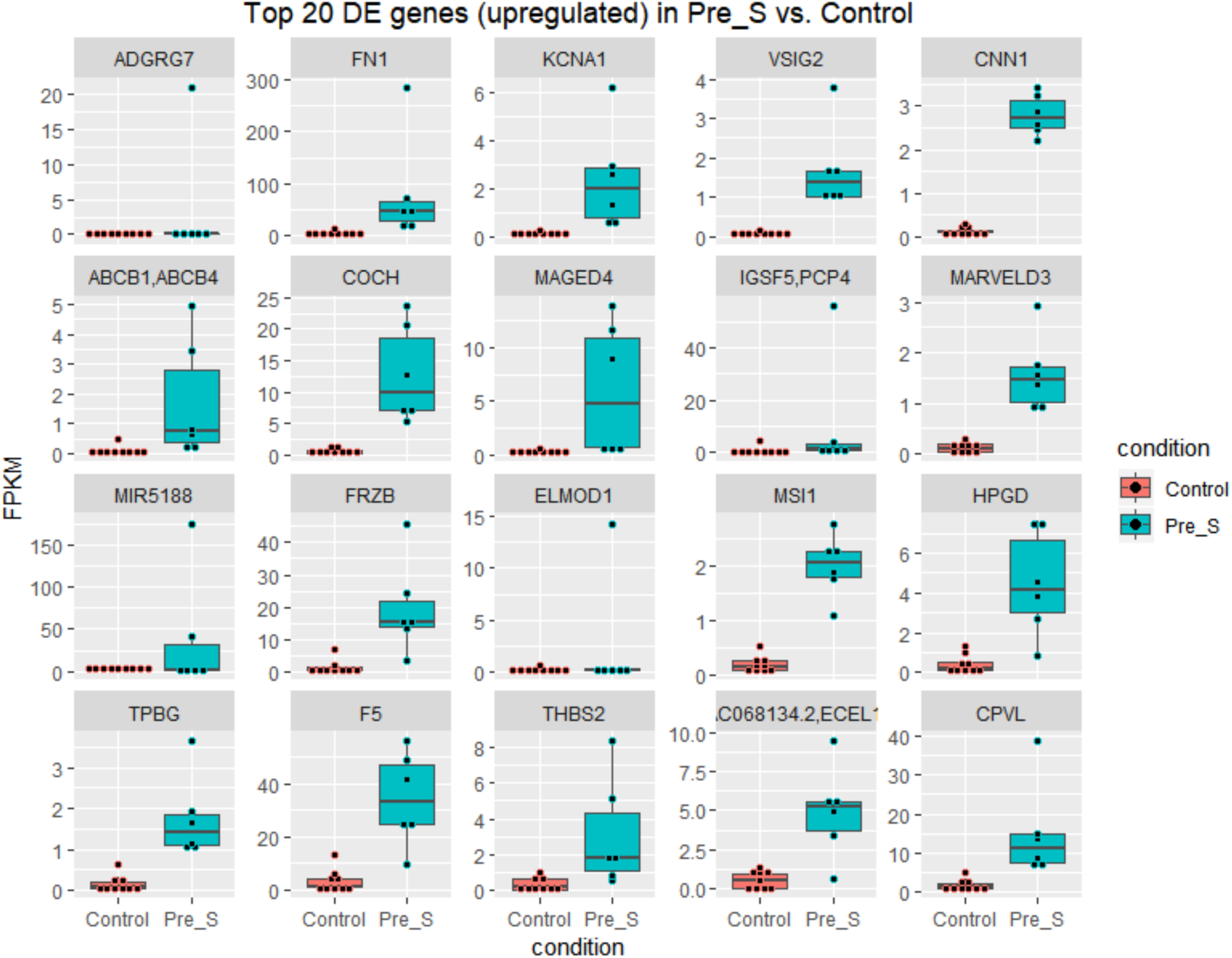
**Top differentially expressed genes in Pre_S tissues**. **(A)** Boxplot showing the level of top 20 upregulated differentially expressed genes in Pre_S tissue replicates.

**Supplemental Figure 4.**
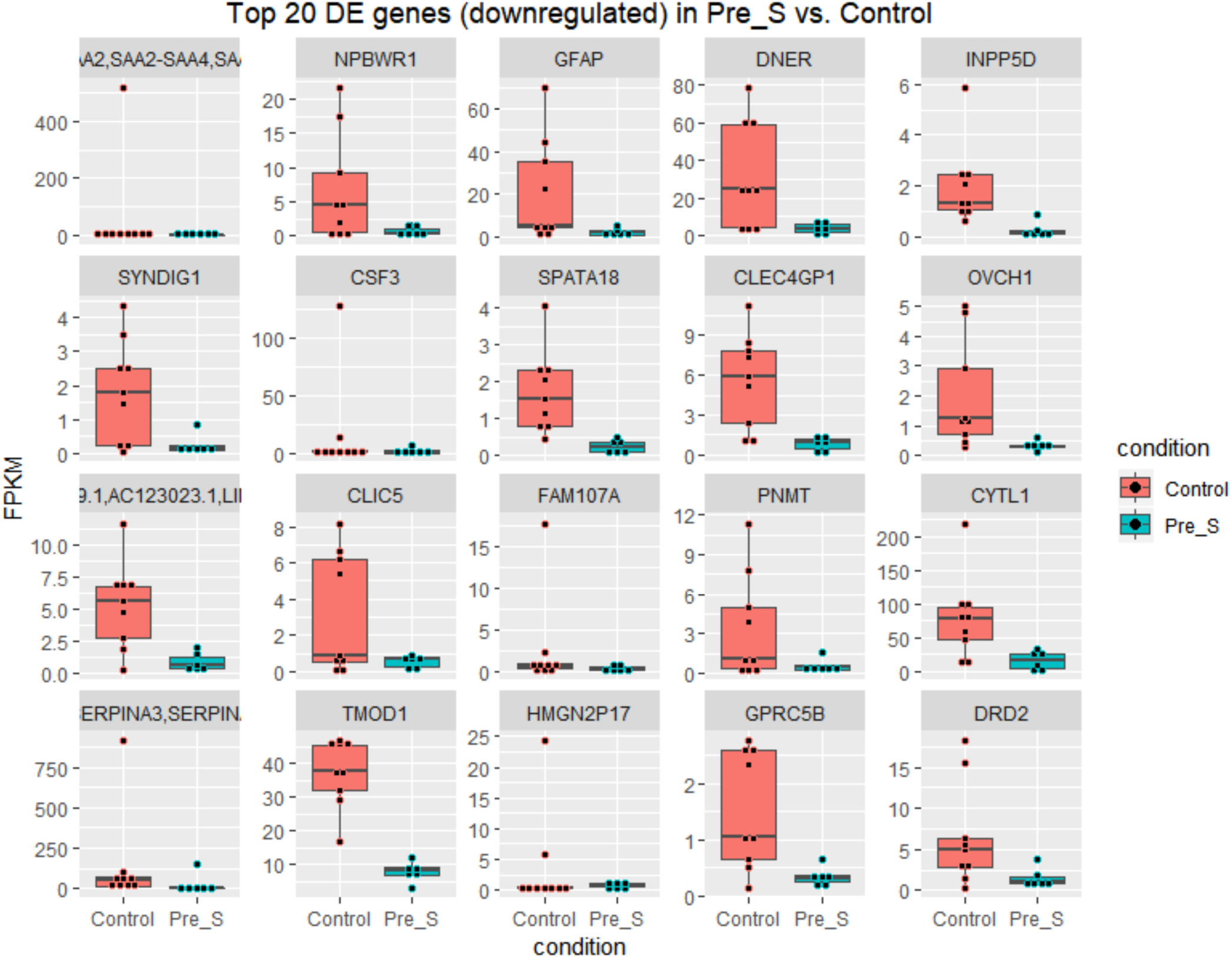
**(B)** Boxplot showing the level of top 20 downregulated differentially expressed genes in Pre_S tissue replicates. A total of 215 differentially expressed genes were identified in Pre_S vs. Control. FPKM > 1.5, fold change > 1.5, FDR <=0.05.

**Supplemental Figure 5.**
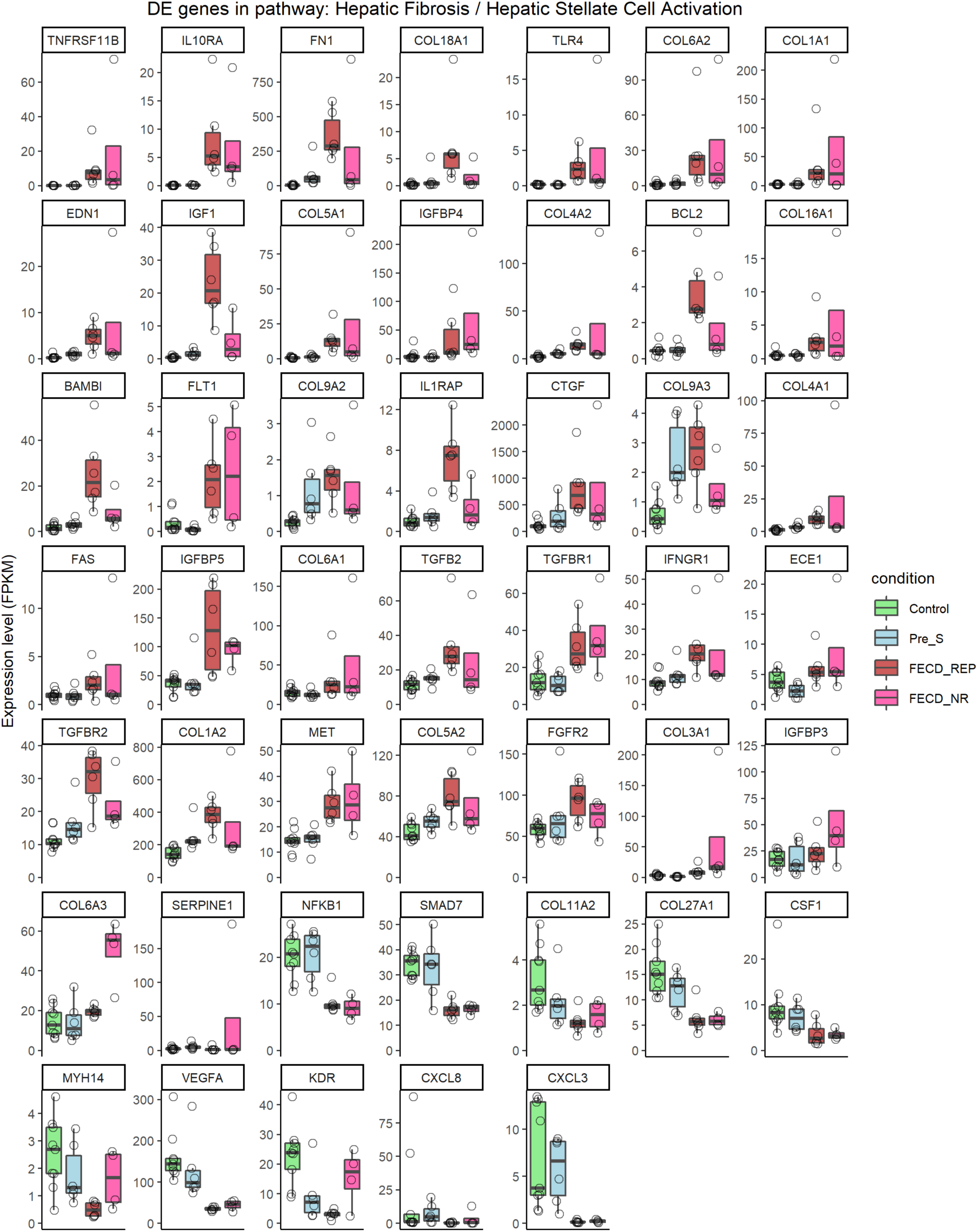
**Detailed pathway analysis. (A).** Expression profile of genes in hepatic fibrosis pathway, all cohorts. FPKM > 1.5, fold change > 1.5, FDR <=0.05.

**Supplemental Figure 5.**
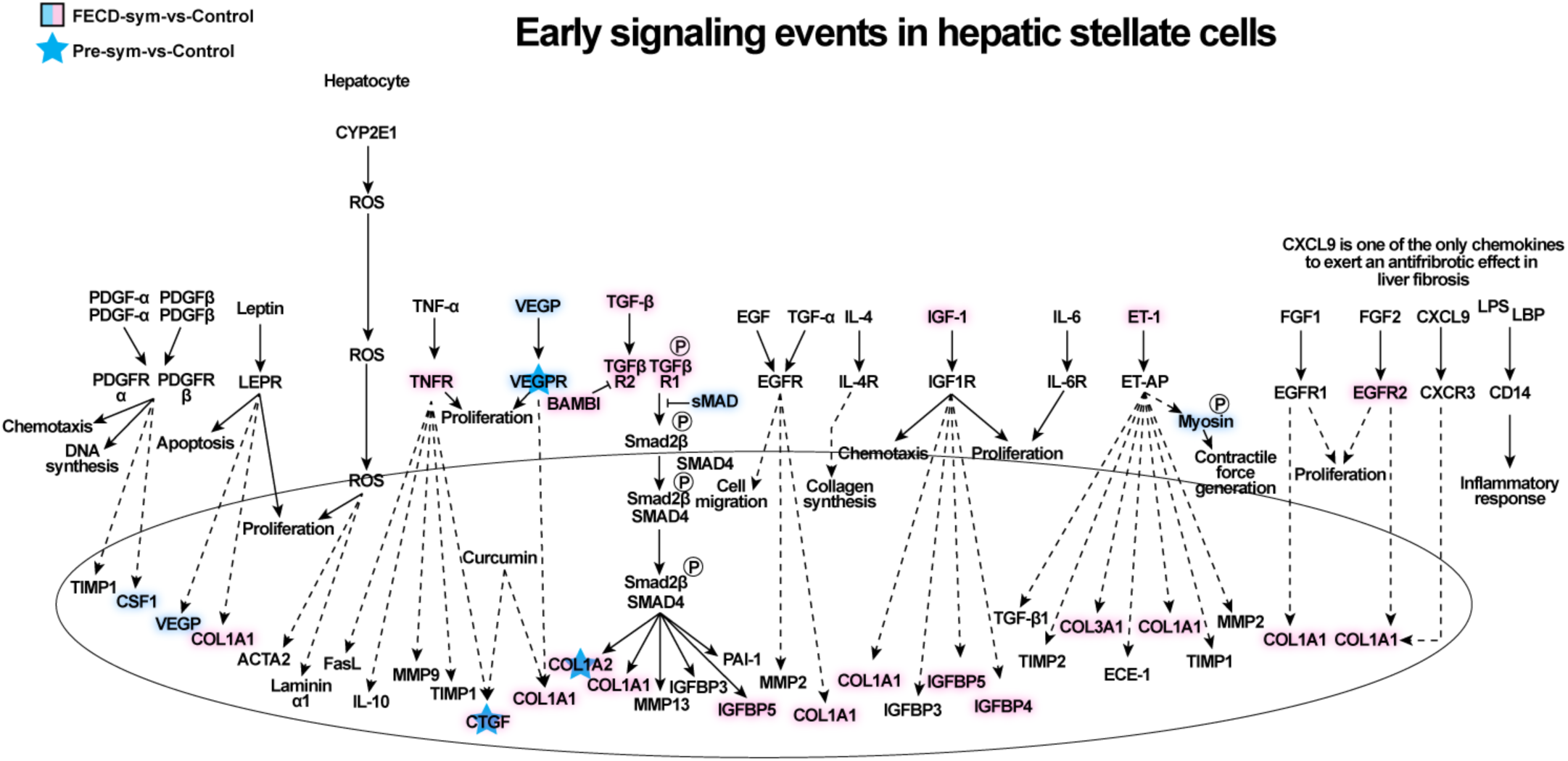
(**B**) Diagram of hepatic fibrosis / hepatic stellate activation pathway based on the comparison of FECD_REP versus Control tissue. Differentially expressed genes that are upregulated are pink and downregulated genes are green. Blue stars mark genes that also change in Pre_S tissue compared to Control samples.

**Supplemental Figure 5.**
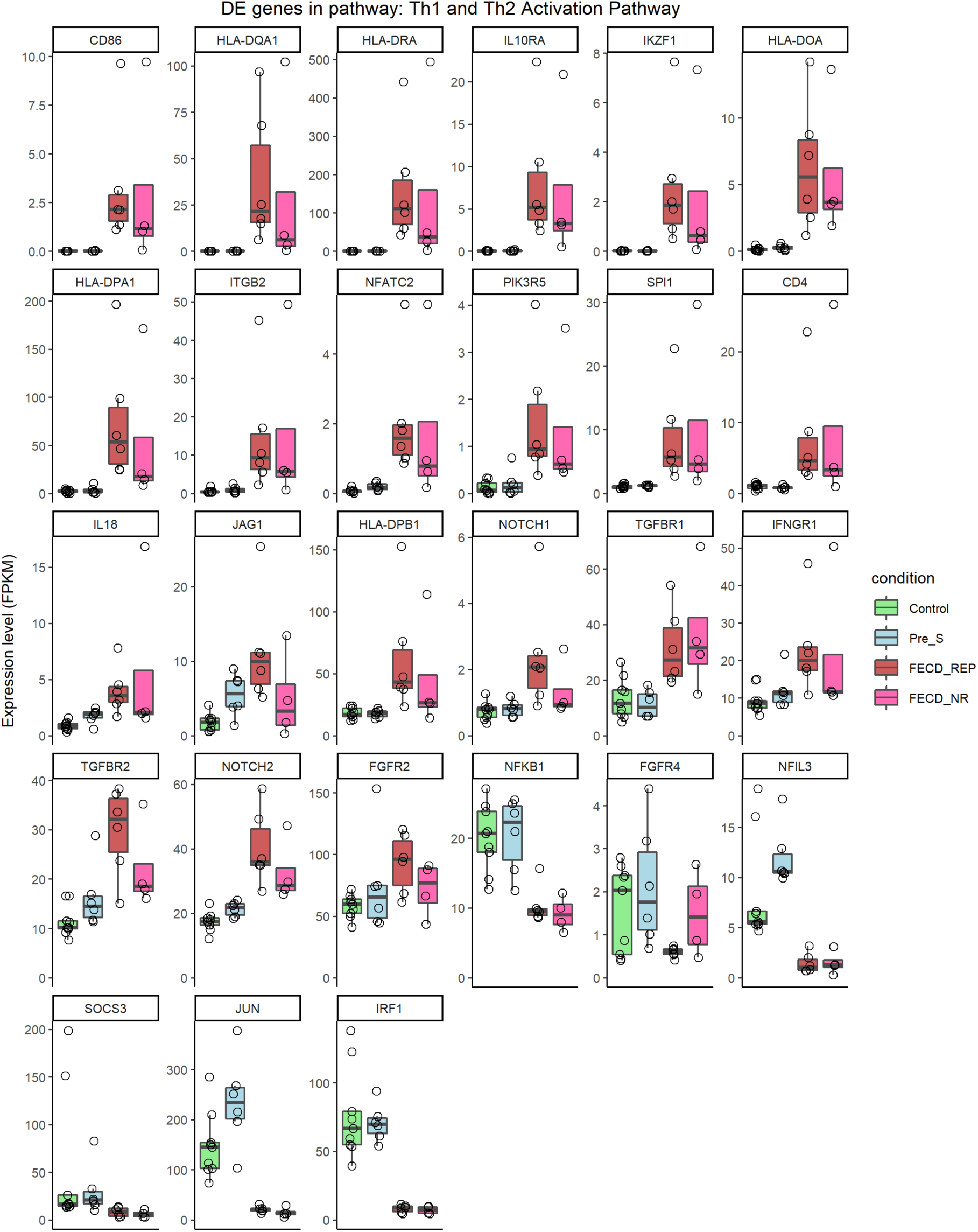
(**C**) Th1/2 activation pathway. FPKM > 1.5, fold change > 1.5, FDR <=0.05.

**Supplemental Figure 5.**
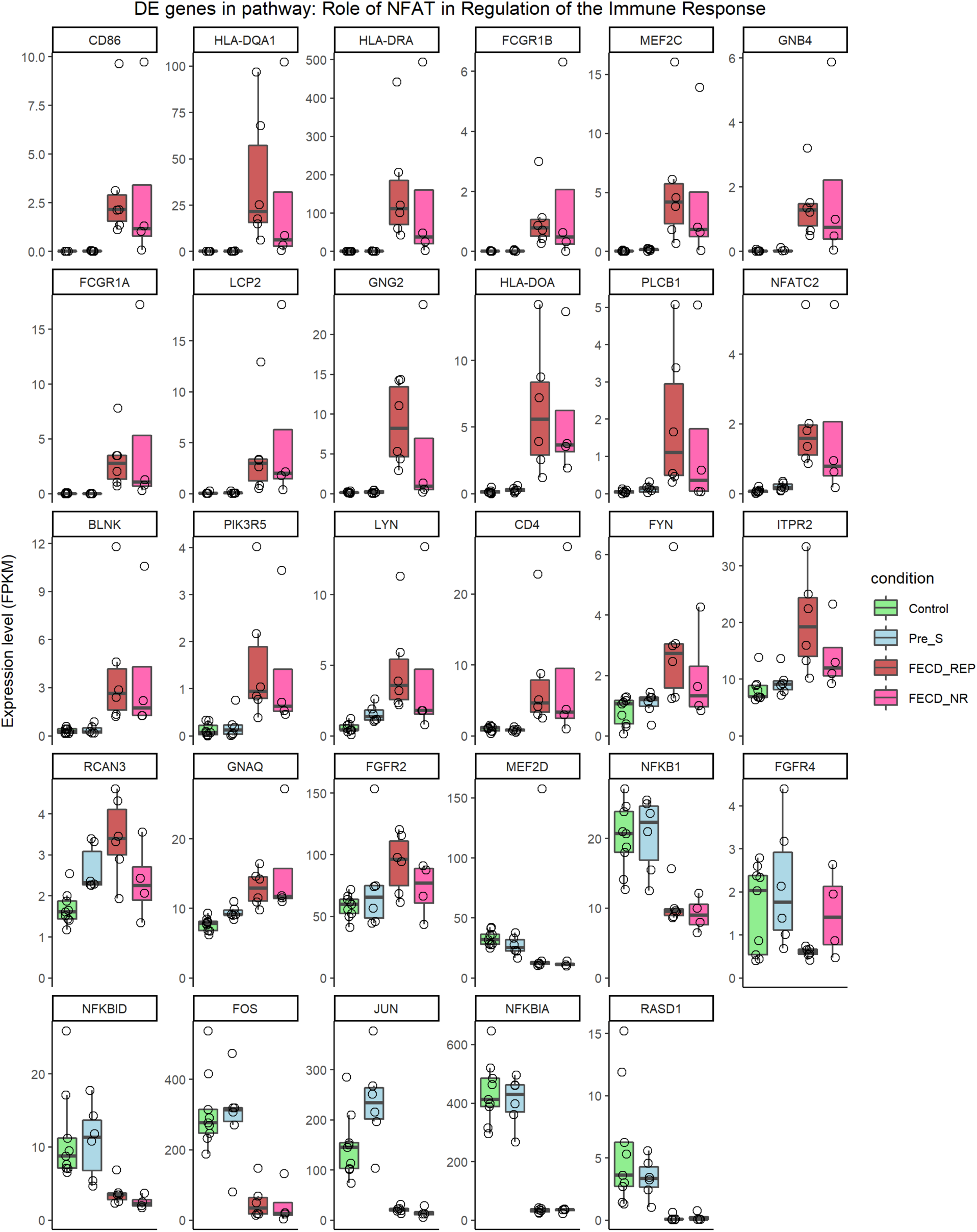
(D) Gene expression changes for NFAT pathway. FPKM > 1.5, fold change > 1.5, FDR <=0.05.

**Supplemental Figure 5.**
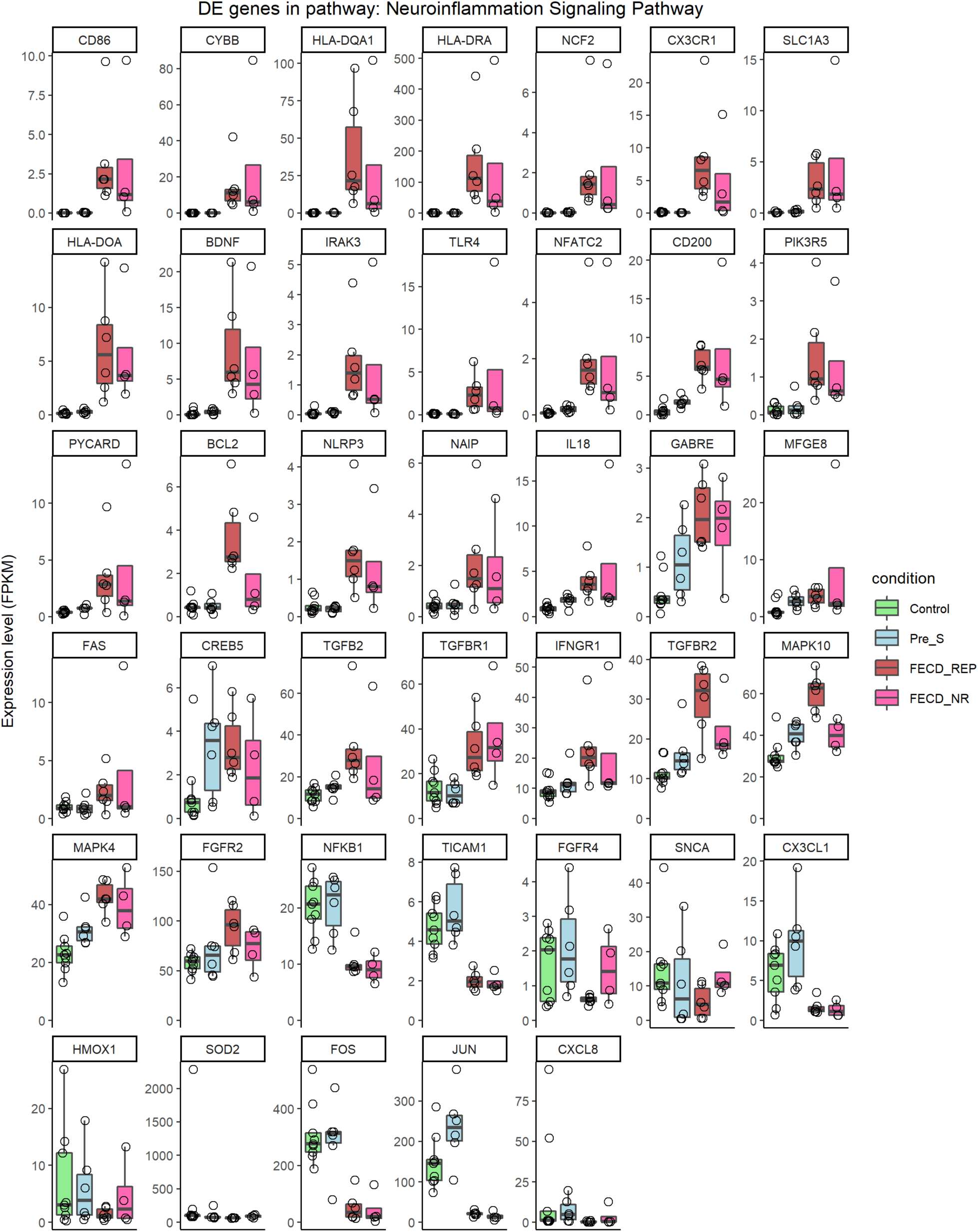
(E) Neuroinflammation Signaling Pathway. FPKM > 1.5, fold change > 1.5, FDR <=0.05.

**Supplemental Figure 5.**
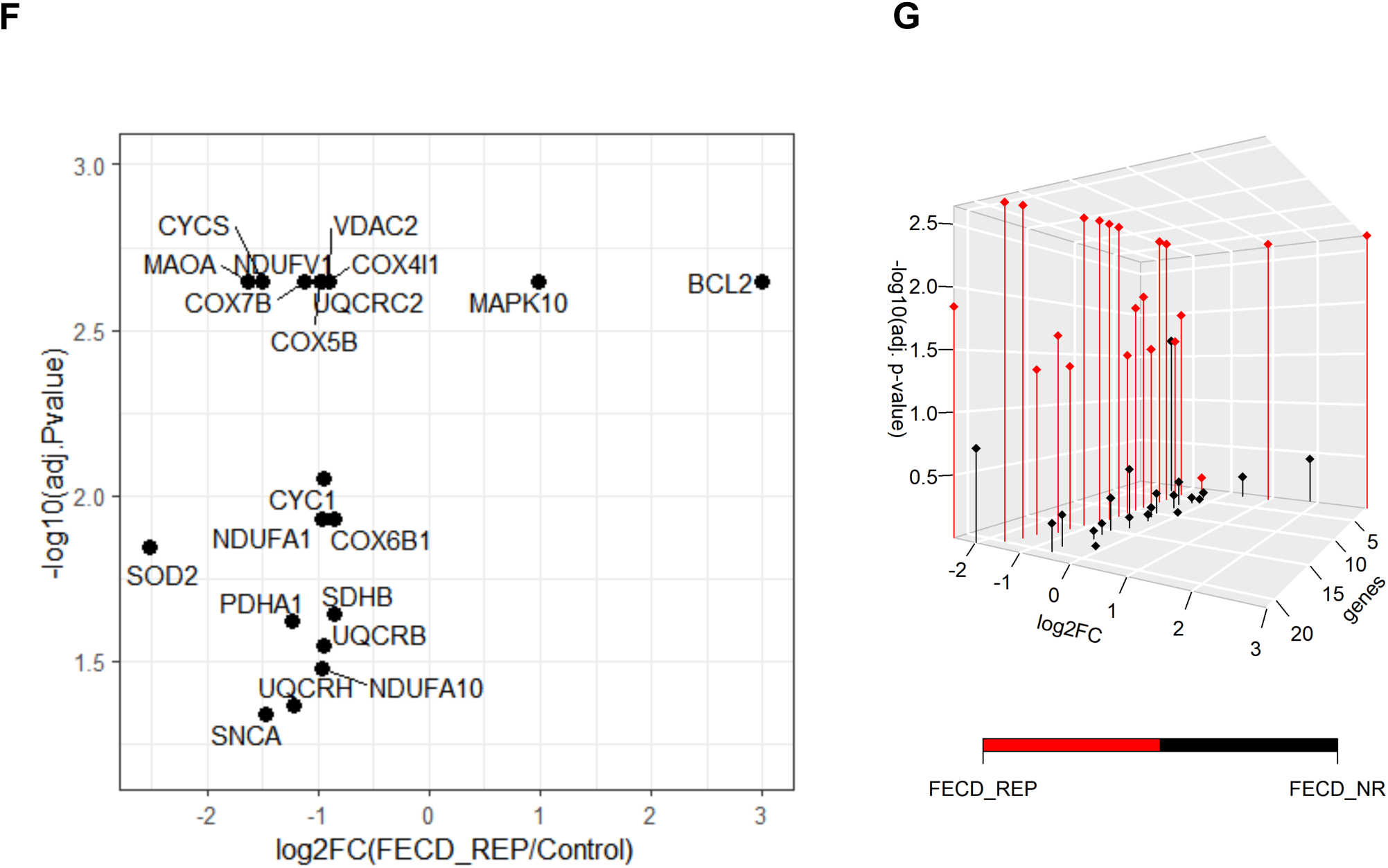
(**F**) Volcano plot of DE genes involved in mitochondrial dysfunction in FECD_REP. (**G**) A scatter 3D plot showing a group of genes involved in mitochondrial dysfunction are differentially expressed in FECD_REP, but not in FECD_NR. X-axis: gene list, y-axis: log2-fold change, z-axis: -log10(adjusted p-value).

**Supplementary Figure 5.**
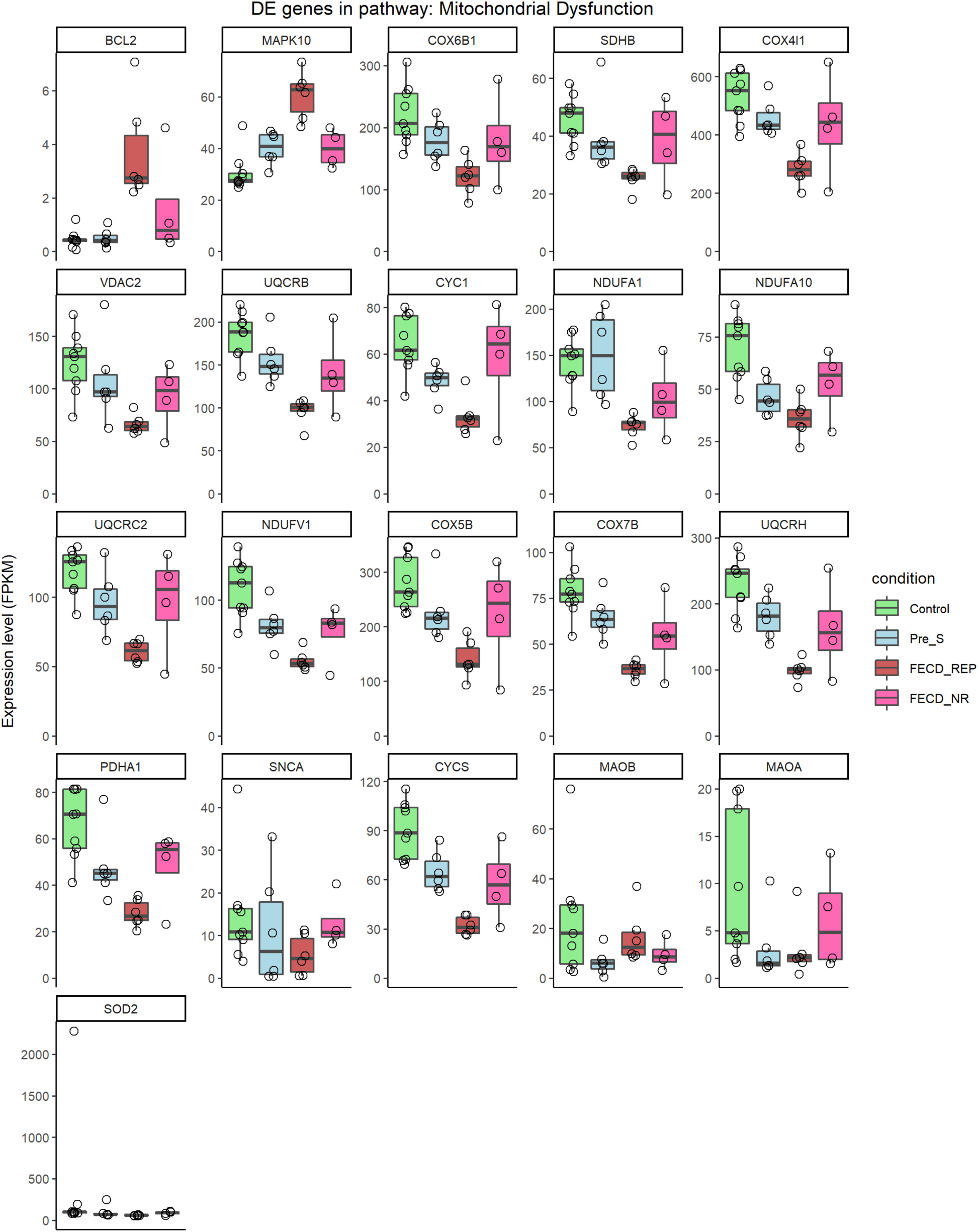
(**H**) Expression changes for mitochondrial genes, all cohorts. FPKM > 1.5, fold change > 1.5, FDR <=0.05.

**Supplemental Table 1.** Characteristics of cornea endothelial tissues used in the experimental validation.

| ID | Age | Phenotype | *TCF4* CUG repeat |
| --- | --- | --- | --- |
| W4056-17-001398 | 54 | Healthy control | 12, 27 |
| W4056-17-001518 | 64 | Healthy control | 14, 26 |
| W4056-17-001449 | 64 | Healthy control | 13, 19 |
| 2015-1477 | 71 | Healthy control | 16, 16 |
| W4056-17-001831 | 59 | Healthy control | 13, 27 |
| W4056-18-002160 | 69 | Healthy control | 15, 15 |
| W4056-17-001754 | 73 | Healthy control | 12, 12 |
| W4056-17-001844 | 66 | Healthy control | 13, 16 |
| W4056-17-002127 | 65 | Healthy control | 12, 18 |
| W4056-17-001239 | 51 | Pre_S | 13, 84 |
| W4056-17-001810 | 57 | Pre_S | 15, 77 |
| W4056-18-002874 | 41 | Pre_S | 17, 86 |
| W4056-19-003507 | 38 | Pre_S | 12, 66 |
| 2015-1615 | 41 | Pre_S | 12, >100 |
| CA208 | 65 | FECD | 18, 60 |
| CA179 | 77 | FECD | 18, 73 |
| CA213 | 67 | FECD | 12, >100 |
| CA219 | 87 | FECD | 18, 76 |
| CA103 | 72 | FECD | 15, 87 |
| CA099 | 70 | FECD | 18, 78 |
| CA072 | 60 | FECD | 16, 84 |
| CA083 | 68 | FECD | 12, 84 |
| CA095 | 91 | FECD | 28, >100 |
| CA209 | 88 | FECD | 15, 53 |
| VVM669 | 66 | FECD | 17, 84 |

**Supplemental Table 2.**
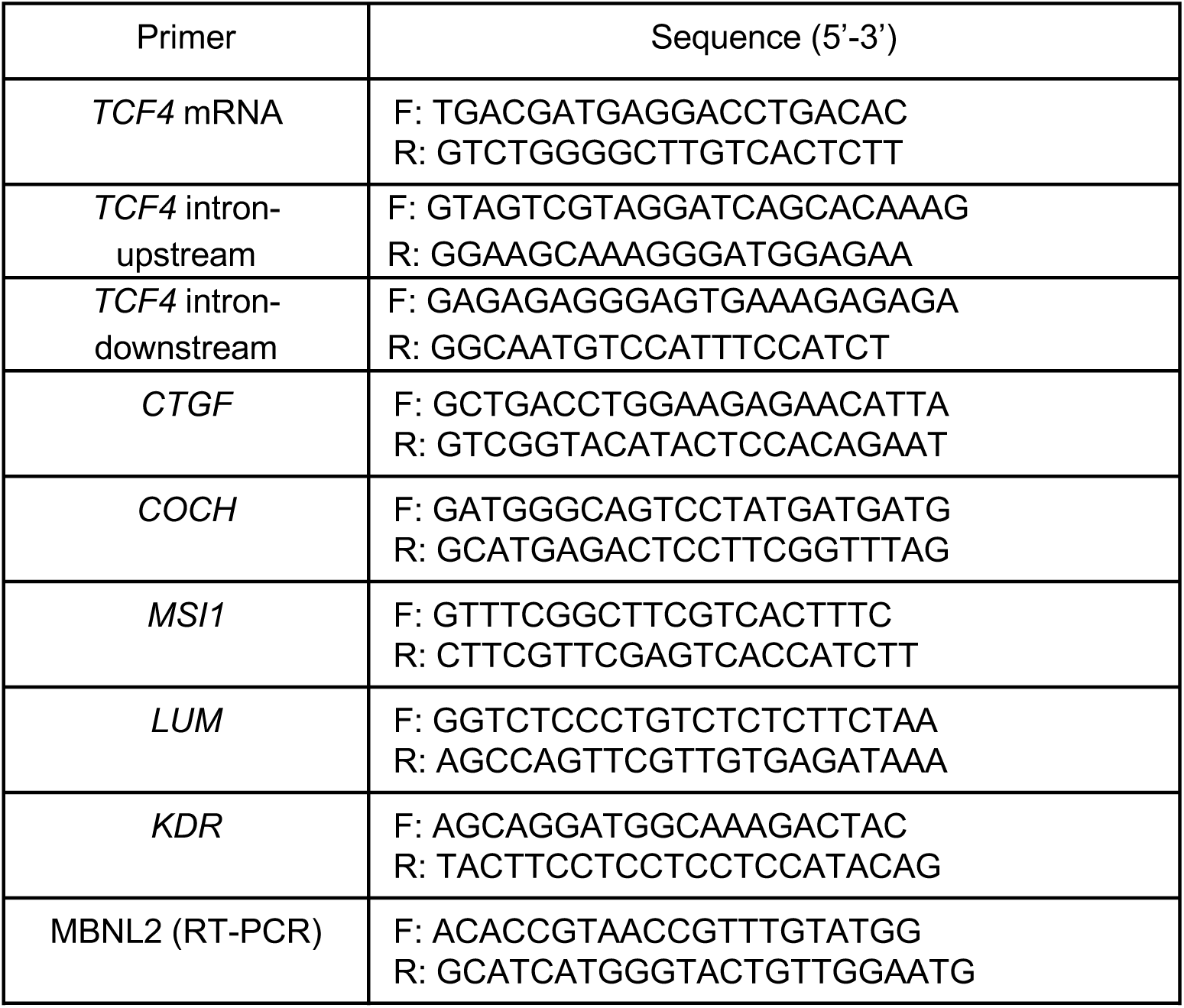
Primer sequences for qPCR or RT-PCR used in the study.

**Supplemental Table 3.**
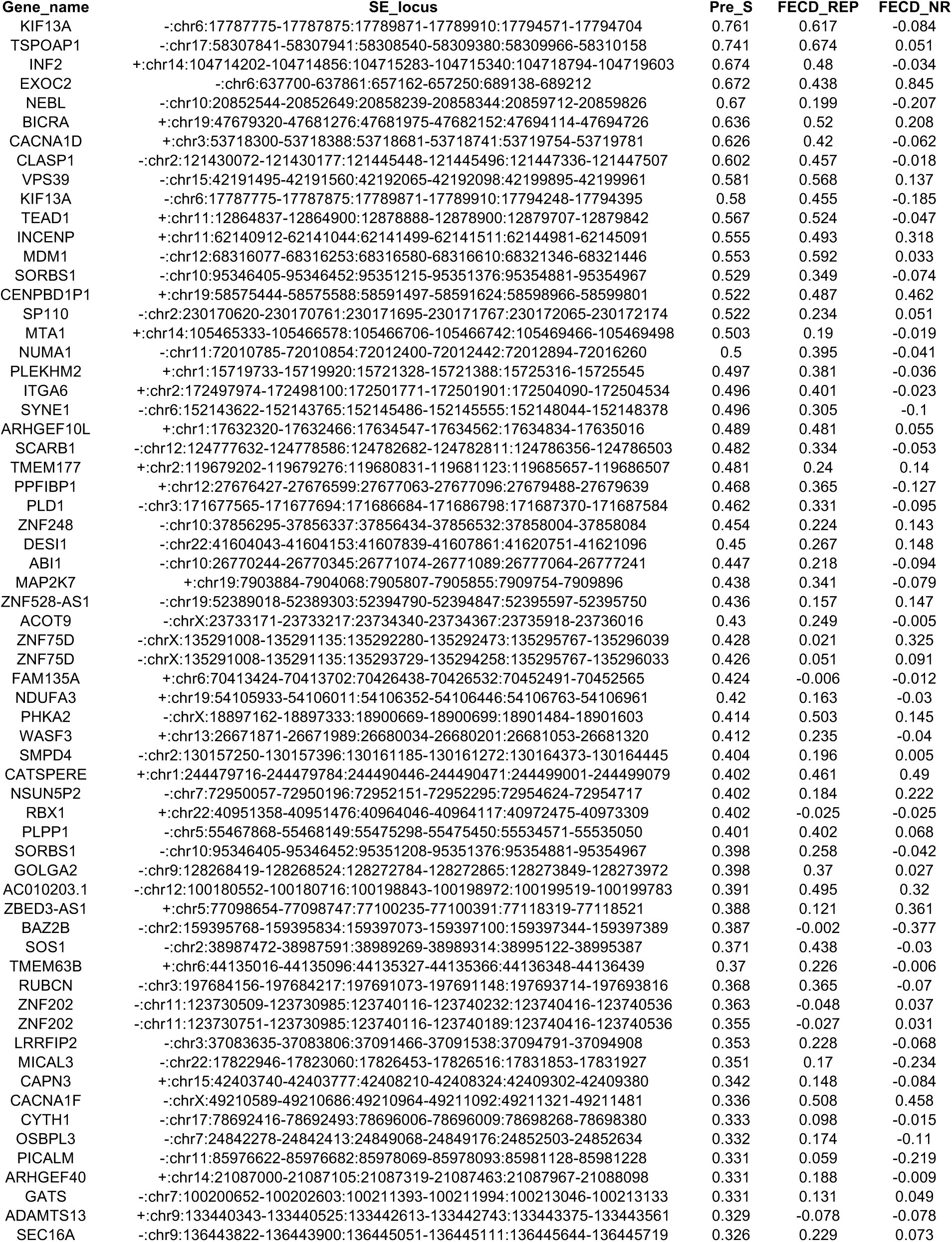
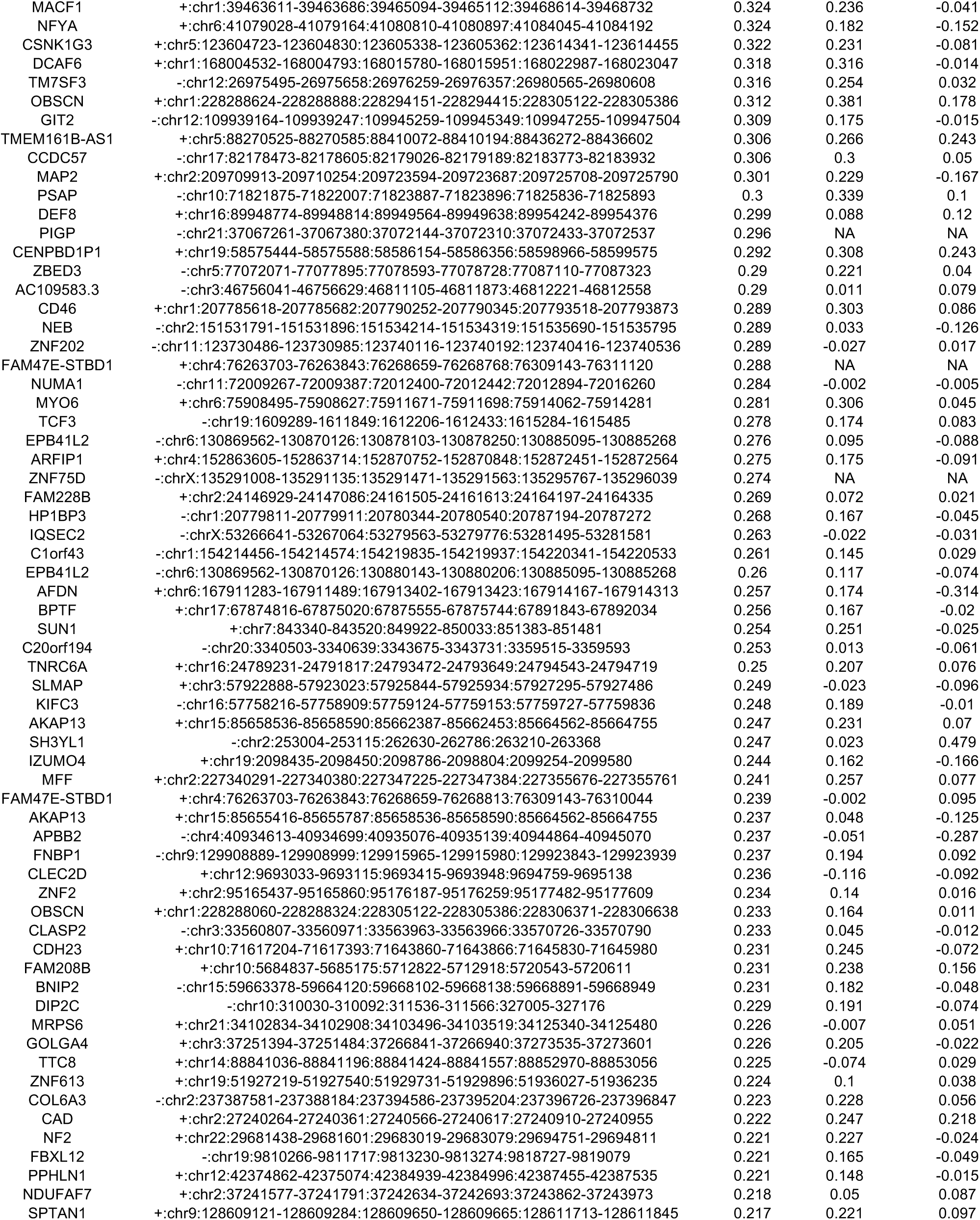
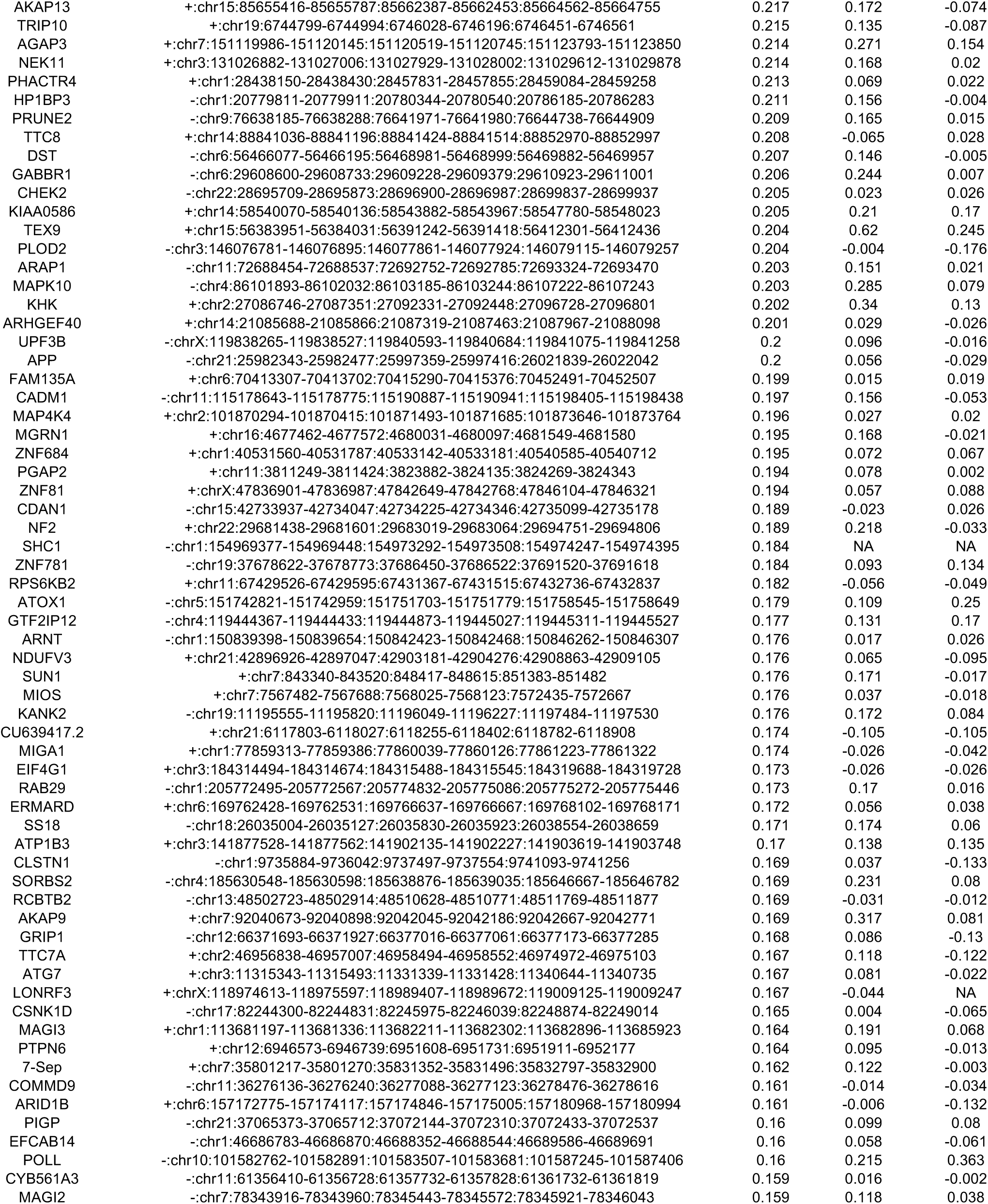
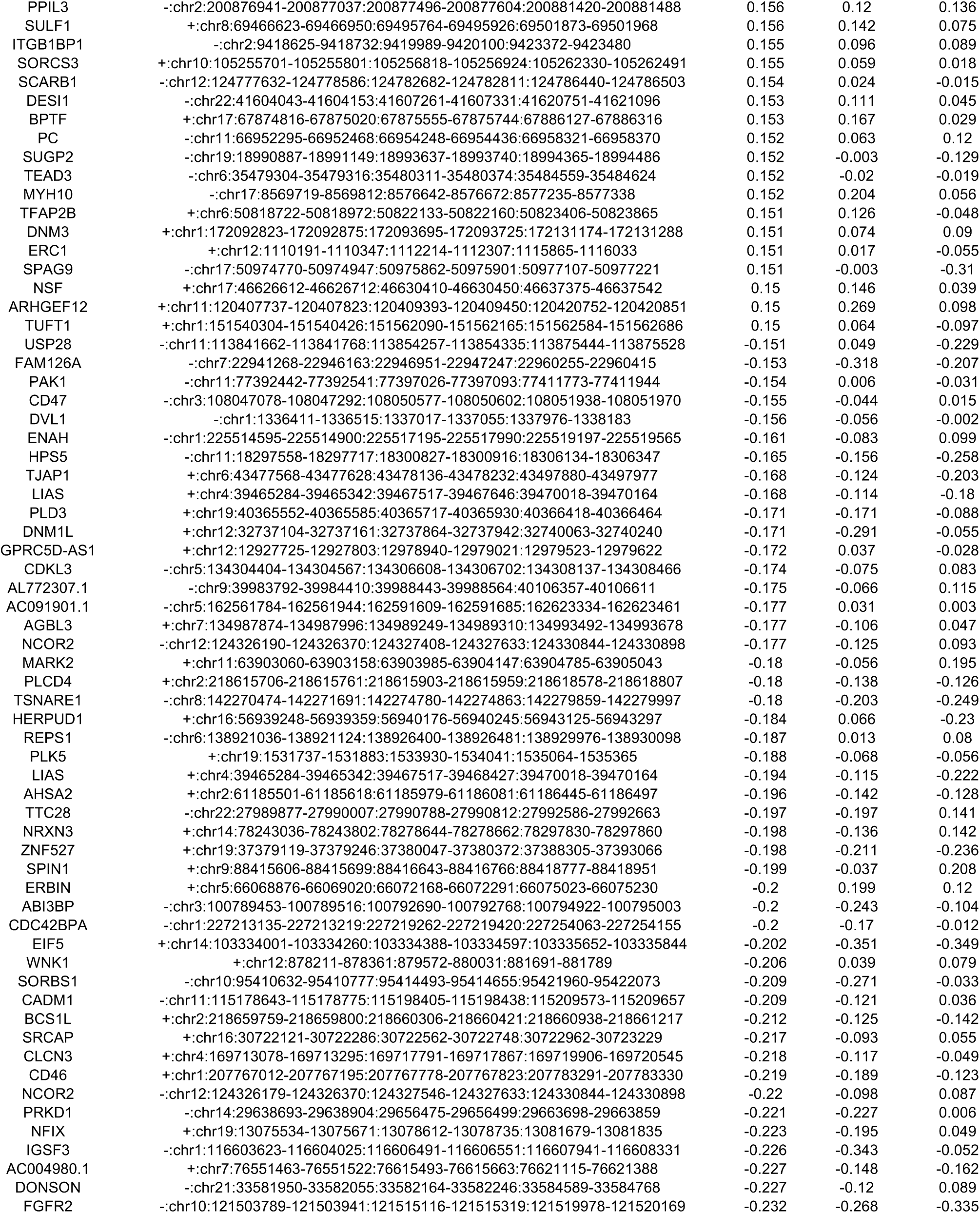
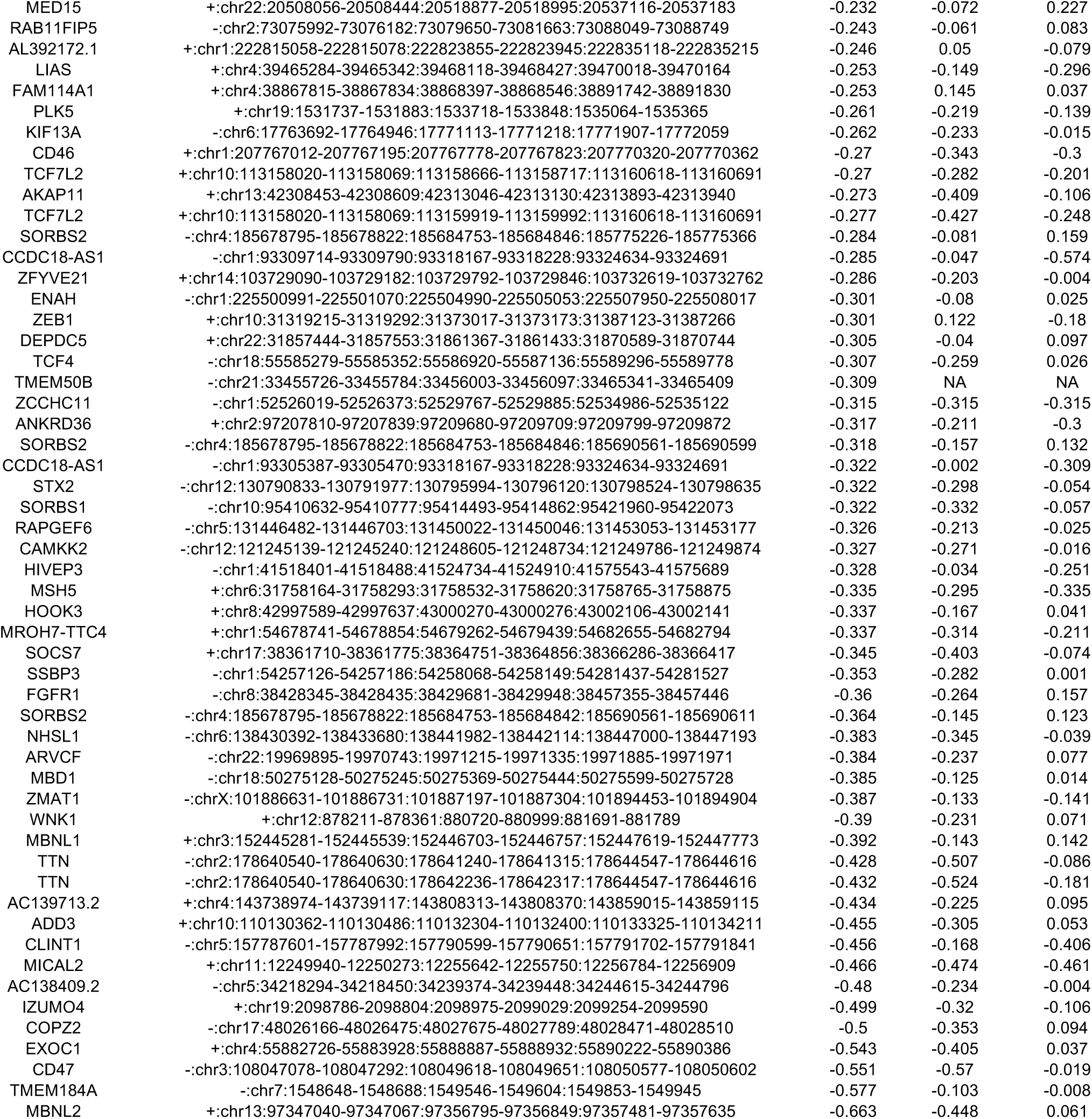
Significant skipped exon events (313) identified in Pre_S. The delta exon inclusion levels for Pre_S, FECD_REP and FECD_NR vs. Control are shown in the last three columns.

**Supplemental Table 4.**
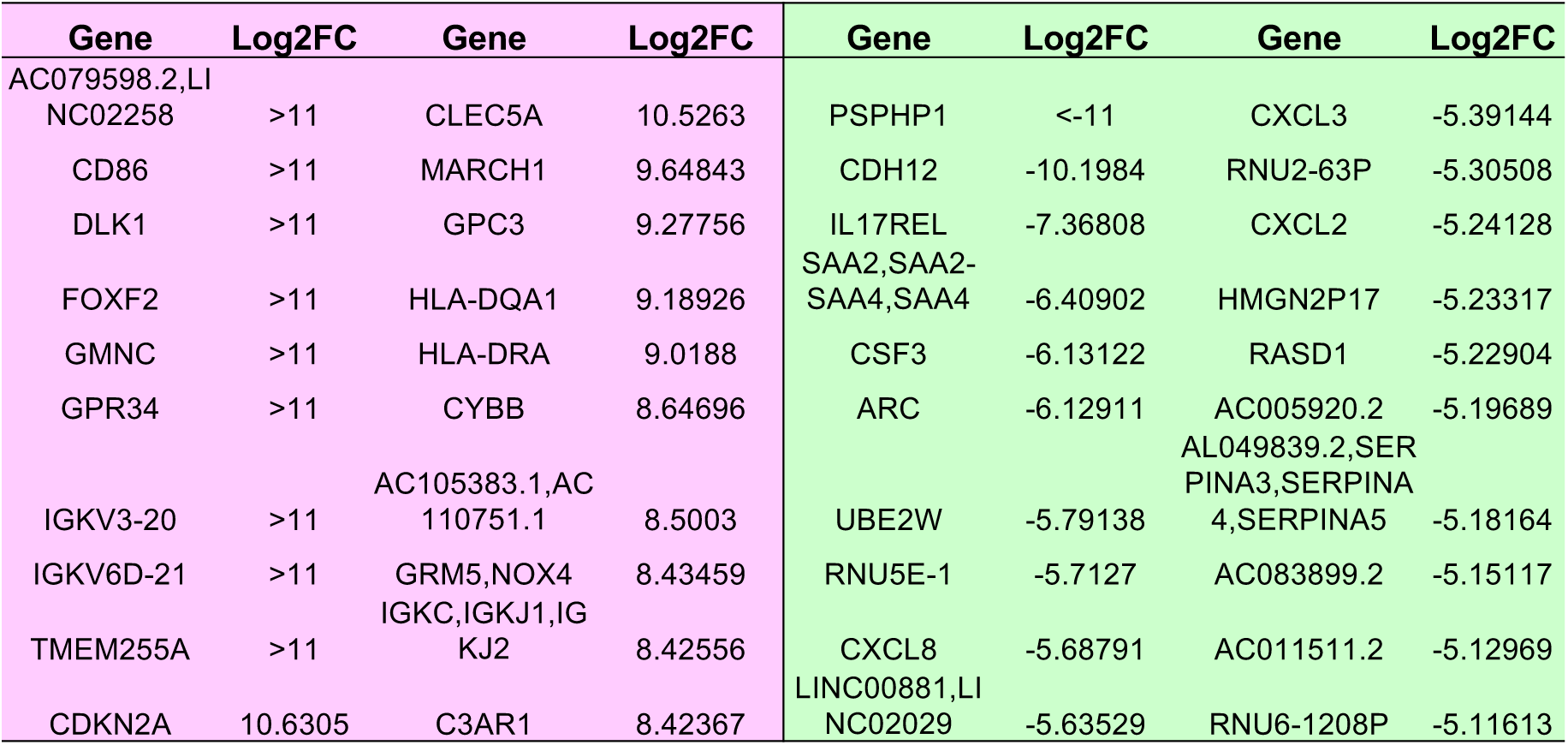
FECD_REP vs. Control top 20 up and down regulated genes.

**Supplemental Table 5.**
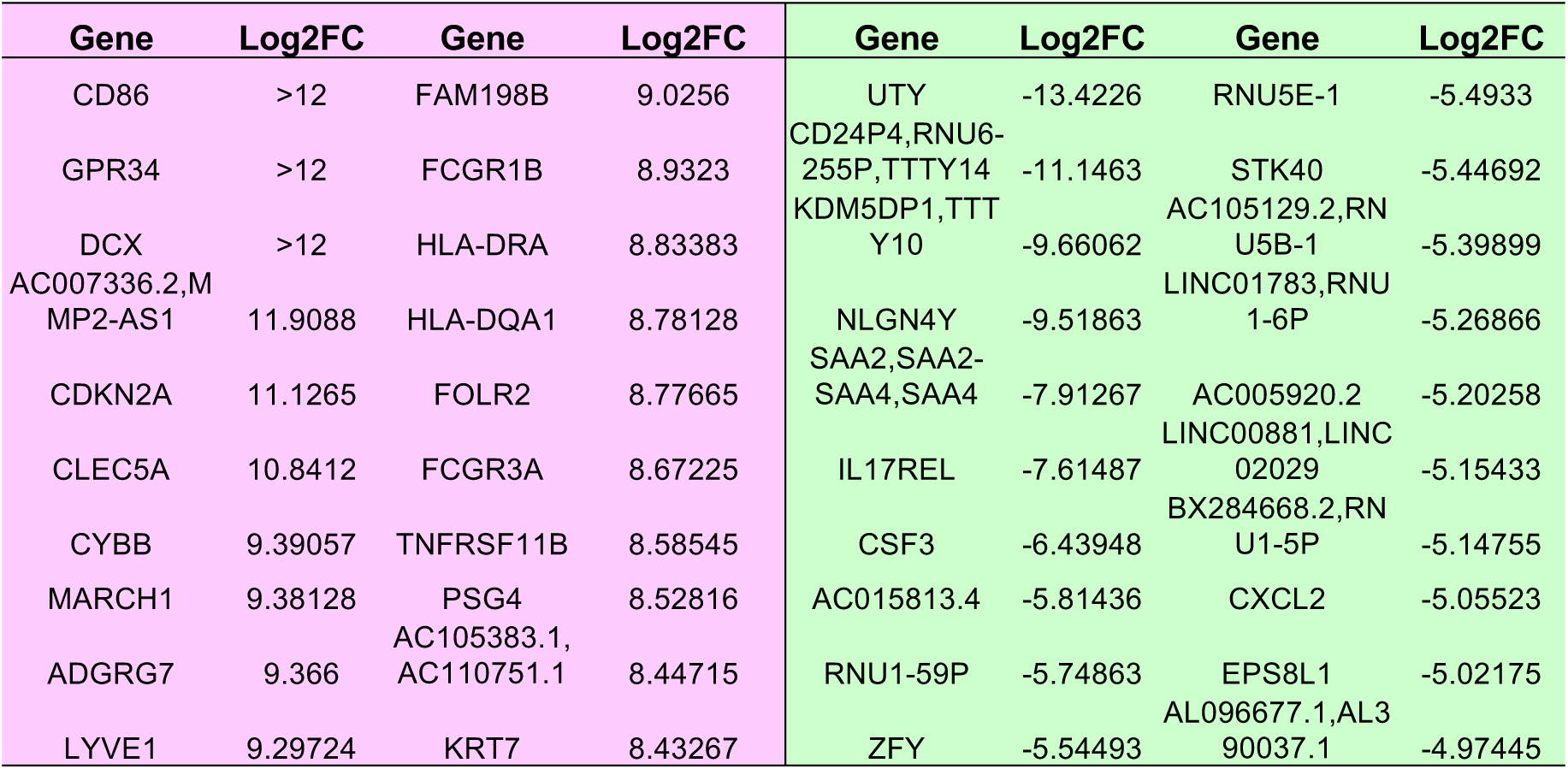
FECD_NR vs. Control top 20 up and down regulated genes.

| Gene | Log2FC | Gene | Log2FC | Gene | Log2FC | Gene | Log2FC |
| --- | --- | --- | --- | --- | --- | --- | --- |
| CD86 | >12 | FAM198B | 9.0256 | UTY | -13.4226 | RNU5E-1 | -5.4933 |
| GPR34 | >12 | FCGR1B | 8.9323 | CD24P4,RNU6-255P,TTTY14 | -11.1463 | STK40 | -5.44692 |
| DCX | >12 | HLA-DRA | 8.83383 | KDM5DP1,TTTY10 | -9.66062 | AC105129.2,RNU5B-1 | -5.39899 |
| AC007336.2,MMP2-AS1 | 11.9088 | HLA-DQA1 | 8.78128 | NLGN4Y | -9.51863 | LINC01783,RNU1-6P | -5.26866 |
| CDKN2A | 11.1265 | FOLR2 | 8.77665 | SAA2,SAA2-SAA4,SAA4 | -7.91267 | AC005920.2 | -5.20258 |
| CLEC5A | 10.8412 | FCGR3A | 8.67225 | IL17REL | -7.61487 | LINC00881,LINC02029 | -5.15433 |
| CYBB | 9.39057 | TNFRSF11B | 8.58545 | CSF3 | -6.43948 | BX284668.2,RNU1-5P | -5.14755 |
| MARCH1 | 9.38128 | PSG4 | 8.52816 | AC015813.4 | -5.81436 | CXCL2 | -5.05523 |
| ADGRG7 | 9.366 | AC105383.1,AC110751.1 | 8.44715 | RNU1-59P | -5.74863 | EPS8L1 | -5.02175 |
| LYVE1 | 9.29724 | KRT7 | 8.43267 | ZFY | -5.54493 | AL096677.1,AL390037.1 | -4.97445 |

**Supplemental Table 6.**
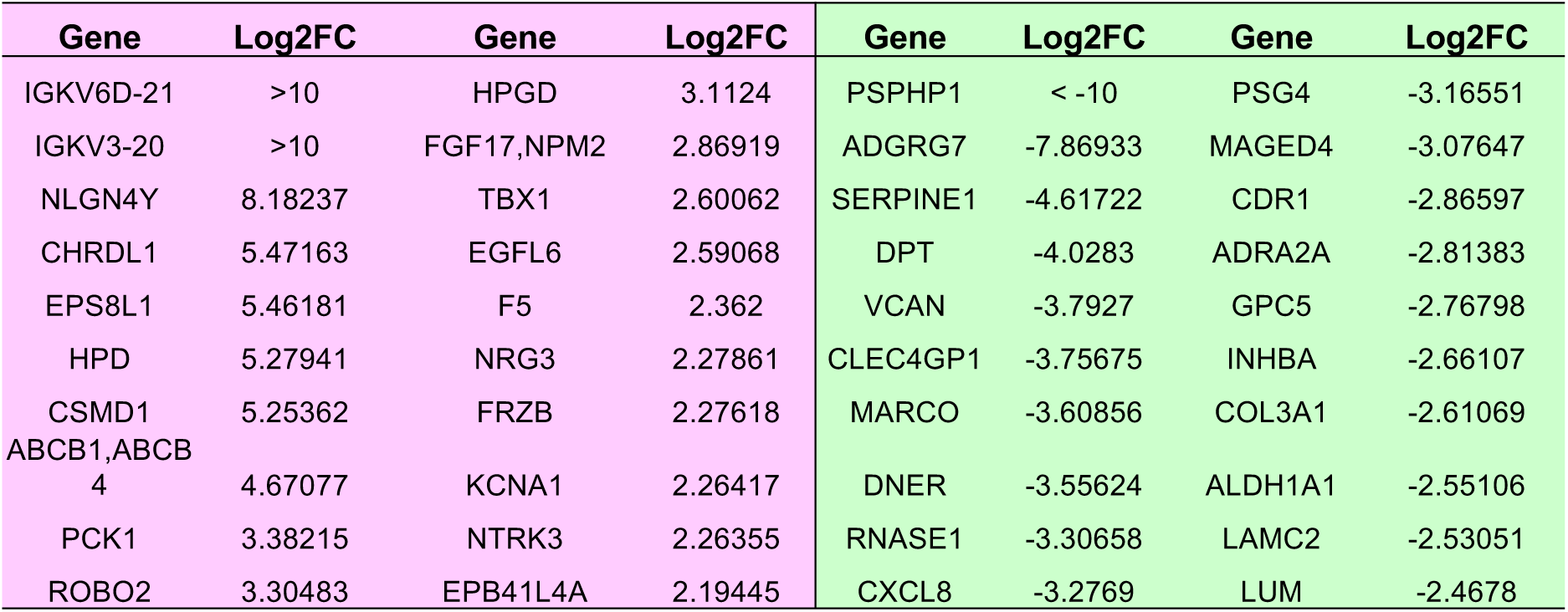
FECD_REP vs. FECD_NR top 20 up and down regulated genes.

**Supplementary Table 7.** The log2FC values and adjusted p-values for 42 fibrosis-pathway associated genes in FECD_REP and Pre_S. These genes were found to be differentially expressed in FECD_REP. For a gene to be considered expressed, its expression level has to be higher than 1.5 FPKM. NA in the following table denotes that a gene is not expressed or its level < 1.5 FPKM.

| Symbol | log2FC_RECD_REP_vs_C | q_value_FECD_REP | log2FC_Pre_S_vs_C | q_value_Pre_S |
| --- | --- | --- | --- | --- |
| COL1A2 | 1.378 | 0.00226 | 0.773381 | 0.00738683 |
| COL3A1 | 1.63 | 0.00226 | -1.71016 | 0.00414755 |
| COL4A2 | 3.045 | 0.00226 | 1.68119 | 0.00226223 |
| COL9A3 | 2.182 | 0.00226 | 2.04259 | 0.00226223 |
| CTGF | 2.723 | 0.00226 | 1.22263 | 0.00226223 |
| FN1 | 7.225 | 0.00226 | 5.08031 | 0.00226223 |
| KDR | -2.927 | 0.00226 | -1.30521 | 0.00226223 |
| BAMBI | 3.936 | 0.00226 | 0.880549 | 0.703482 |
| COL11A2 | -1.337 | 0.0363 | -0.534815 | 0.824853 |
| COL1A1 | 4.067 | 0.00226 | 0.17908 | 0.999784 |
| COL27A1 | -1.302 | 0.00226 | -0.406276 | 0.549341 |
| COL4A1 | 3.112 | 0.00226 | 1.70879 | 0.149814 |
| COL5A2 | 0.829 | 0.00226 | 0.294152 | 0.70792 |
| COL6A1 | 1.075 | 0.00891 | -0.215987 | 0.999784 |
| COL6A2 | 4.794 | 0.00226 | 0.90987 | 0.750228 |
| CSF1 | -1.53 | 0.00226 | -0.496574 | 0.51737 |
| CXCL3 | -5.391 | 0.00226 | -0.288417 | 0.999784 |
| CXCL8 | -5.688 | 0.00226 | -1.30287 | 0.156507 |
| FGFR2 | 0.681 | 0.0457 | 0.369273 | 0.658643 |
| IFNGR1 | 1.289 | 0.00226 | 0.355356 | 0.868174 |
| IGFBP4 | 2.646 | 0.00226 | -1.01873 | 0.542776 |
| IGFBP5 | 1.89 | 0.00226 | 0.336963 | 0.756909 |
| IL1RAP | 2.861 | 0.00226 | 0.716579 | 0.486386 |
| MET | 1.009 | 0.00226 | 0.0417758 | 0.999784 |
| MYH14 | -2.326 | 0.00226 | -0.504027 | 0.802993 |
| NFKB1 | -0.957 | 0.0457 | 0.0288995 | 0.999784 |
| SMAD7 | -1.079 | 0.0169 | -0.048409 | 0.999784 |
| TGFB2 | 1.616 | 0.00226 | 0.418369 | 0.659669 |
| TGFBR1 | 1.235 | 0.00415 | -0.249032 | 0.999784 |
| TGFBR2 | 1.378 | 0.00226 | 0.50825 | 0.396355 |
| VEGFA | -2.232 | 0.00226 | -0.326395 | 0.74898 |
| BCL2 | 3.004 | 0.00226 | NA | NA |
| COL16A1 | 2.454 | 0.00226 | NA | NA |
| COL18A1 | 4.992 | 0.00226 | NA | NA |
| COL5A1 | 4.67 | 0.00226 | NA | NA |
| COL9A2 | 2.557 | 0.00226 | NA | NA |
| EDN1 | 4.109 | 0.00226 | NA | NA |
| FLT1 | 2.443 | 0.00415 | NA | NA |
| IGF1 | 6.021 | 0.00226 | NA | NA |
| IL10RA | 7.616 | 0.00226 | NA | NA |
| TLR4 | 4.397 | 0.00226 | NA | NA |
| TNFRSF11B | 7.586 | 0.00226 | NA | NA |

